# Site-specific analysis reveals candidate cross-kingdom small RNAs, tRNA and rRNA fragments, and signs of fungal RNA phasing in the barley-powdery mildew interaction

**DOI:** 10.1101/2022.07.26.501657

**Authors:** Stefan Kusch, Mansi Singh, Hannah Thieron, Pietro D. Spanu, Ralph Panstruga

**Affiliations:** Unit of Plant Molecular Cell Biology, Institute for Biology I, RWTH Aachen University, Worringerweg 1, D-52056 Aachen, Germany; Imperial College, London, United Kingdom

**Keywords:** RNA interference, powdery mildew, barley, *Blumeria*, haustorium, mycelium, extracellular vesicles, cross-kingdom RNAi, tRNA fragments, rRNA fragments, phased RNA, small RNA

## Abstract

The establishment of host-microbe interactions requires molecular communication between both partners, which involves the mutual transfer of noncoding small RNAs. Previous evidence suggests that this is also true for the barley powdery mildew disease, which is caused by the fungal pathogen *Blumeria hordei*. However, previous studies lacked spatial resolution regarding the accumulation of small RNAs upon host infection by *B. hordei*. Here, we analysed site-specific small RNA repertoires in the context of the barley-*B. hordei* interaction. To this end, we dissected infected leaves into separate fractions representing different sites that are key to the pathogenic process: epiphytic fungal mycelium, infected plant epidermis, isolated haustoria, a vesicle-enriched fraction from infected epidermis, and extracellular vesicles. Unexpectedly, we discovered enrichment of specific 31- to 33-base long 5’-terminal fragments of barley 5.8S ribosomal RNA (rRNA) in extracellular vesicles and infected epidermis, as well as particular *B. hordei* tRNA fragments in haustoria. We describe canonical small RNAs from both the plant host and the fungal pathogen that may confer cross-kingdom RNA interference activity. Interestingly, we found first evidence of phased small RNAs (phasiRNAs) in *B. hordei*, a feature usually attributed to plants, which may be associated with the post-transcriptional control of fungal coding genes, pseudogenes, and transposable elements. Our data suggests a key and possibly site-specific role for cross-kingdom RNA interference and noncoding RNA fragments in the host-pathogen communication between *B. hordei* and its host barley.

## Introduction

All complex multicellular organisms, including plants, require the exchange of information between cells for development and reproduction, but also need to communicate signals between each other and to coordinate the response to external stimuli. This exchange is referred to as either intra- or inter-organismal communication, depending on whether it takes place between cells of the same organism or between separate organisms. A special case of inter-organismal communication is the exchange of information between organisms of different taxa. Examples of this phenomenon in the area of plant-microbe interactions comprise mutually advantageous symbioses and diseases caused by pathogenic fungi (1, 2). Gene regulation by non-coding small RNAs (sRNAs) can play an important role in the context of such plant-microbe encounters (1).

In plants, sRNAs are important for regulating and fine-tuning diverse processes including growth and development, maintenance of genome integrity, epigenetic inheritance, and facilitating responses to both abiotic and biotic stress (3, 4). Based on their biogenesis and function, sRNAs can be divided into two major subclasses, micro RNAs (miRNAs) and small-interfering RNAs (siRNAs, (5)). *Micro RNA* (*MIR*) genes encode pri-miRNAs, which are further processed into pre-miRNAs and finally mature miRNAs. In eukaryotes other than mammals and plants, such as in fungi, it is challenging to prove the presence of *bona fide* miRNAs due to lack of evidence for the non-functional miRNA precursor strand; for this reason, they are termed miRNA-like RNAs (milRNAs; (6)). In contrast to miRNAs, siRNAs are generated from double-stranded DNA molecules of various origins (5). In some cases, miRNAs trigger the generation of secondary siRNAs, for example phased siRNAs (phasiRNAs; i.e., regularly spaced siRNAs that derive from a common precursor RNA), which are sometimes associated with the silencing of transposable elements (7). If the sRNA that triggers the formation of phasiRNAs is derived from a remote *trans*-acting sRNA locus, it is termed *trans*-acting siRNA (tasiRNA; (7)). While phasiRNAs and tasiRNAs are well-described in plants, they are largely unexplored in fungi, as we are aware of only one study reporting their existence in the fungal kingdom (8).

Both classes of sRNAs can induce either transcriptional gene silencing or posttranscriptional gene silencing of specific target genes, collectively referred to as RNA interference (RNAi; (9)). Irrespective of the class, biogenesis and function of the sRNA, RNAi involves cleavage of a double-stranded RNA (dsRNA) molecule by a dicer-like (Dcl) ribonuclease and loading of the mature sRNA onto an Argonaute (Ago) protein, together forming the RNA-induced silencing complex (5). Dcl proteins process the dsRNA precursor fragments into smaller pieces of 21-24 nucleotides or bases. The length of the fragment produced depends on the specific Dcl proteins, which recognise different types of dsRNA. The fragments are then bound by Ago proteins that specifically distinguish different sRNA lengths and are associated with distinct modes of gene silencing. For example, *Arabidopsis thaliana* AGO1 binds 21 base-long miRNAs generated by DCL1 or DCL4 and executes posttranscriptional gene silencing by cleaving target mRNAs in a sequence-specific manner. By contrast, *A. thaliana* AGO4 binds almost exclusively 24 base-long siRNAs generated by DCL3. AGO4 then induces transcriptional gene silencing by triggering DNA methylation, a process referred to as RNA-dependent DNA methylation (10). In each case, the Ago-bound sRNA determines target specificity through sequence complementarity and the Ago protein is the key effector for gene silencing. In the context of inter-organismal communication, the phenomenon of sRNA-triggered trans-species gene silencing has been termed cross-kingdom RNAi. One of the best studied systems for plant-microbe cross-kingdom RNAi is the *A. thaliana-Botrytis cinerea* pathosystem, where exchange of sRNAs occurs in both directions (11, 12). Once fungal sRNAs reach plant cells they are loaded onto *A. thaliana* Ago proteins, which carry out the cleavage of host target transcripts such as those encoding mitogen-activated protein kinases (11). Similarly, the most abundant *A. thaliana* sRNAs detected in *B. cinerea* cells originate from tasiRNAs or intergenic loci, and target fungal genes important for pathogenicity (13).

Extracellular vesicles (EVs) may serve as shuttles for inter- and intraorganismal RNA molecule transfer in the context of plant-microbe interactions (13). The release of EVs and their contents into the plant apoplast upon pathogen challenge and their delivery to infection sites was first described over 50 years ago (14, 15). This parallels the situation in mammalian cells where EVs in the intercellular space target nearby and distant cells and are important components of intra-organismal communication (16). The EVs interact with recipient cells to deliver cargo like sRNAs by membrane fusion and vesicle internalisation (17–19). EVs occur in all pro- and eukaryotic phyla (20). However, it is still unclear whether EV shuttling is the main mechanism for the delivery of sRNAs in plant-pathogen interactions, since evidence for the shuttling of sRNAs outside of EVs was recently found in *A. thaliana* (21).

The agronomically important crop barley (*Hordeum vulgare*) is the host plant of the ascomycete fungus *Blumeria hordei*, previously named *B. graminis* f.sp. *hordei* (22), which causes the powdery mildew disease on barley. *B. hordei* colonises barley biotrophically and the obligate relationship between host and pathogen requires tightly controlled gene regulation. In a previous study, we found that the essential components of the RNAi machinery are present in *B. hordei* (23). Moreover, we discovered at least 1,250 sRNA loci in the *B. hordei* genome, of which 524 were predicted to have mRNA targets in the host barley. Expression of sRNAs in both barley and *B. hordei* is consistent with a role in regulating transcript abundance in both partners during the interaction (24). In *B. hordei*, milRNAs may control the mRNA levels of fungal virulence genes whereas barley miRNAs and phasiRNAs might control transcript levels of components of the plant innate immune system. Inter-organismal exchange of RNAs is thought to be mediated by membrane-bound vesicles (13). Interestingly, vesicle-like structures have been observed at the interface between *B. hordei* infection structures and barley leaf epidermal cells (25, 26); this finding would be consistent with an exchange of sRNAs mediated by EVs in the context of the barley-powdery mildew interaction.

In this study, we analysed the sRNA spectrum in the *B. hordei* - barley pathosystem. To this end, we dissected infected leaves into separate fractions representing different sites that are key to the pathogenic process, i.e., epiphytic fungal mycelium, infected plant epidermis, isolated haustoria, a vesicle-enriched fraction from infected epidermis, and EVs. Unexpectedly, we discovered enrichment of specific 31- to 33-base long 5’ fragments of barley 5.8S ribosomal RNA (rRNA) in EVs, vesicles, and infected epidermis, as well as specific *B. hordei* tRNA fragments in haustoria. We describe canonical sRNAs from both the plant host and the fungal pathogen that may confer cross-kingdom RNAi activity. Interestingly, we found first evidence of phasiRNAs in *B. hordei*, a feature usually attributed to plants, which may be associated with the post-transcriptional control of coding genes, pseudogenes, and transposable elements. Our data suggests a key role for cross-kingdom RNAi and noncoding RNA fragments in the host-pathogen communication between *B. hordei* and its host barley.

## Results

### Site-specific sRNA sampling

Based on three independent experiments, we isolated total RNA from the following biological materials (“sites”) of *B. hordei*-infected barley leaves four days after inoculation (Figure 1, see Materials and Methods for a detailed description of the samples): (1) Epiphytic fungal mycelium (MYC), (2) infected epidermis without mycelium (EPI), (3) fungal haustoria (HAU), (4) microsomes of the epidermis without mycelium (P40). In addition, RNA from (5) apoplastic extracellular vesicles (EV+) were isolated from *B. hordei*-infected barley leaves three days after inoculation. As a control, we also isolated (6) total RNA from extracellular vesicles of non-infected plants (EV-). The 18 RNA samples (6 sources, 3 replicates each) were then used to extract RNA, which was subsequently subjected to Illumina-based short read sequencing at a depth of 30 million (MYC, EPI, HAU, P40) or 20 million (EV+, EV-) reads per sample (Supplementary Table 1).

**Figure 1.**
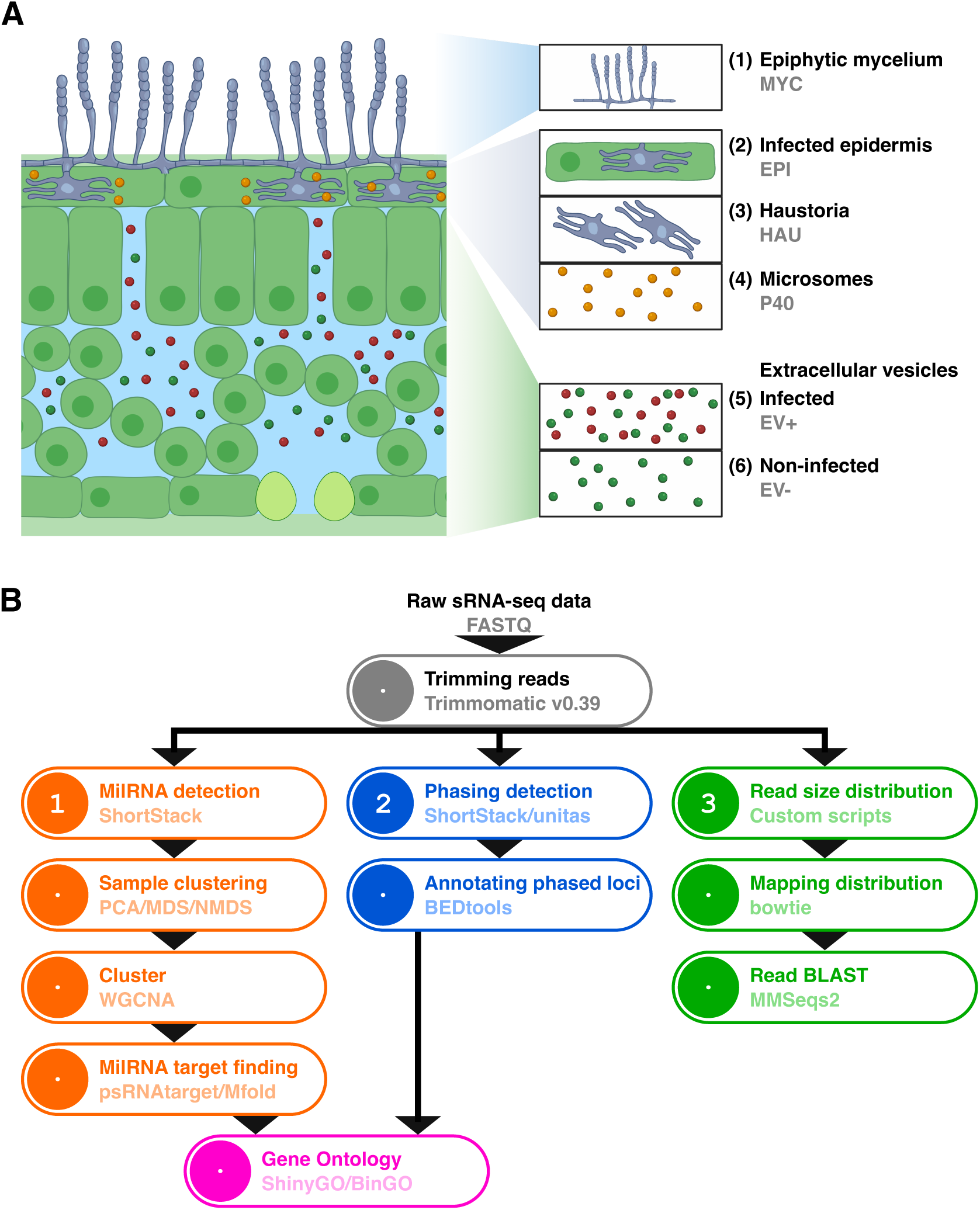
Isolation of total RNA from six distinct sample types. (**A**) We isolated total RNA from the following biological materials of *B. hordei*-infected barley leaves at four days after inoculation: (1) Epiphytic fungal mycelium (MYC), (2) infected epidermis without mycelium (EPI), (3) fungal haustoria (HAU), (4) microsomes of the epidermis depleted of haustoria (P40). In addition, we isolated total RNA from (5) apoplastic extracellular vesicles of infected plants (EV+) at three days after inoculation, and (6) apoplastic extracellular vesicles from non-infected control plants (EV-). The figure was created using bioRender.com. (**B**) Bioinformatic pipeline for sRNA-seq data analysis. We analysed the sRNA-seq reads in three ways: (1) by read size and read mapping distribution to the respective genomes, followed by read BLAST against the RFAM database for particular fractions of the read data; (2) by ShortStack analysis to detect putative milRNAs in both organisms, followed by principal component analysis for sample clustering, milRNA expression analysis, milRNA target prediction, and functional description of these targets by GO assignment; (3) by detection of loci enriched with predicted phasiRNAs.

### Distinct rRNA- and tRNA-derived sRNAs are enriched in *B. hordei*- and barley-derived sample materials

The length of the trimmed sRNA reads ranged from 15 to 75 bases. However, the majority of the reads were between 15 and 40 bases long; between 282,734 (2.4%) and 5,432,088 (13.8%) per sample cumulatively accounted for reads between 41-74 bases (Supplementary Table 2). We determined the sRNA length distribution profiles from the 15 to 40 base reads of the various samples and found that each of the six biological materials showed a distinctive and largely reproducible pattern: the sRNAs from mycelium (MYC) exhibited a bimodal distribution with a prominent peak at 21 and 22 bases, and a much shallower and broader peak with a maximum at 29-33 bases. Likewise, the epidermal (EPI) samples had a bimodal size distribution with marked peaks at 21-22 and 32-33 bases, respectively. The HAU and P40 samples exhibited a broad size range without any outstanding peak, but an increase in the number of 27-32 bases reads compared to reads below or above this range. Finally, the two EV sample types (EV- and EV+) exhibited distinctive profiles: the EV+ size distribution was reminiscent of EPI and MYC samples, with a bimodal distribution and characteristic peaks at 21-23 and 31-32 bases, respectively. However, the EV-sRNAs showed a single prominent peak at 31-32 bases (Supplementary Figure 1A).

We mapped the sRNA reads to the respective reference genomes of *H. vulgare* IBSCv2 (27) and *B. hordei* DH14 (28) using bowtie within the ShortStack (29) pipeline. Overall, between 3,715,684 and 32,777,442 reads could be assigned to the *H. vulgare* genome and between 431,626 and 22,069,070 reads to the *B. hordei* genome, respectively (Supplementary Table 1). We next determined the number of reads mapping to each of the genomes separately for the read sizes of 15-40 bases (Supplementary Figure 1B; Supplementary Table 3 and 4). We noted that reads below 19 bases could not be unequivocally allocated to either organism by read mapping alone, resulting in >100% total mapping counts. Hence, we considered reads <19 bases as ambiguous and disregarded these from further read length-related analysis. We found that the majority of 21-22 base reads in the MYC and EPI samples derived from the *B. hordei* genome, while reads from the 31-32/32-33 base peaks in the EPI and EV samples almost exclusively aligned to the genome of *H. vulgare*. The majority of reads >19 bases from the P40 and EV-samples were from the *H. vulgare* genome, while reads from the EPI, HAU, and EV+ samples originated from both organisms. Regardless of the read length, except for the distinct 31/32/33 base peak in reads from *H. vulgare*, the majority of the reads originated from either coding genes or transposable elements (Supplementary Figure 2).

We next explored the identity and possible molecular origin of the sRNAs from the very prominent 31-32 base peak in the EV+ and 32-33 base peak in the EPI samples, both from *B. hordei*-infected barley leaves. Using MMSeqs2 for BLAST against the RNA families (RFAM) databases of conserved noncoding RNAs (i.e., transfer RNAs, ribosomal RNAs, small nucleolar RNAs, small nuclear RNAs), we found that >80% of the sequences in these two sample types corresponded to a specific short fragment of the barley 5.8S rRNA (Figure 2). This 5.8S rRNA fragment (rRF) was specifically found in 31-32-base long reads in EV+ and EV−, and 32-33-base long reads in EPI (Supplementary Figure 3). In case of the EV-sample, approximately 30% of 31-32 base reads match this barley 5.8S rRF, while only <15% of P40 and HAU reads originate from it. Between 15 and 30% of the reads in the MYC and HAU samples were identified as derived from *B. hordei* 5.8S rRNA (Figure 2). In HAU, *B. hordei* 5.8S rRNA fragments were abundant in reads of 27-32 bases in length (Supplementary Figure 3). Notably, the vast majority of the 5.8S rRNA-associated reads mapped to the 3’-end of the molecule in both organisms, even in samples lacking a distinctive peak at 31-33 bases (>90%; Figure 2B and Table 1). We refer to these regions as *H. vulgare* rRNA fragment *Hvu*-rRF0001 and *B. hordei* rRNA fragment *Bho*-rRF0001 (Figure 2B and Table 2). In case of *B. hordei*, we found a second less abundant fragment adjacent to *Bho*-rRF0001, called *Bho*-rRF0002 (Supplementary Figure 3 and Table 2).

**Figure 2.**
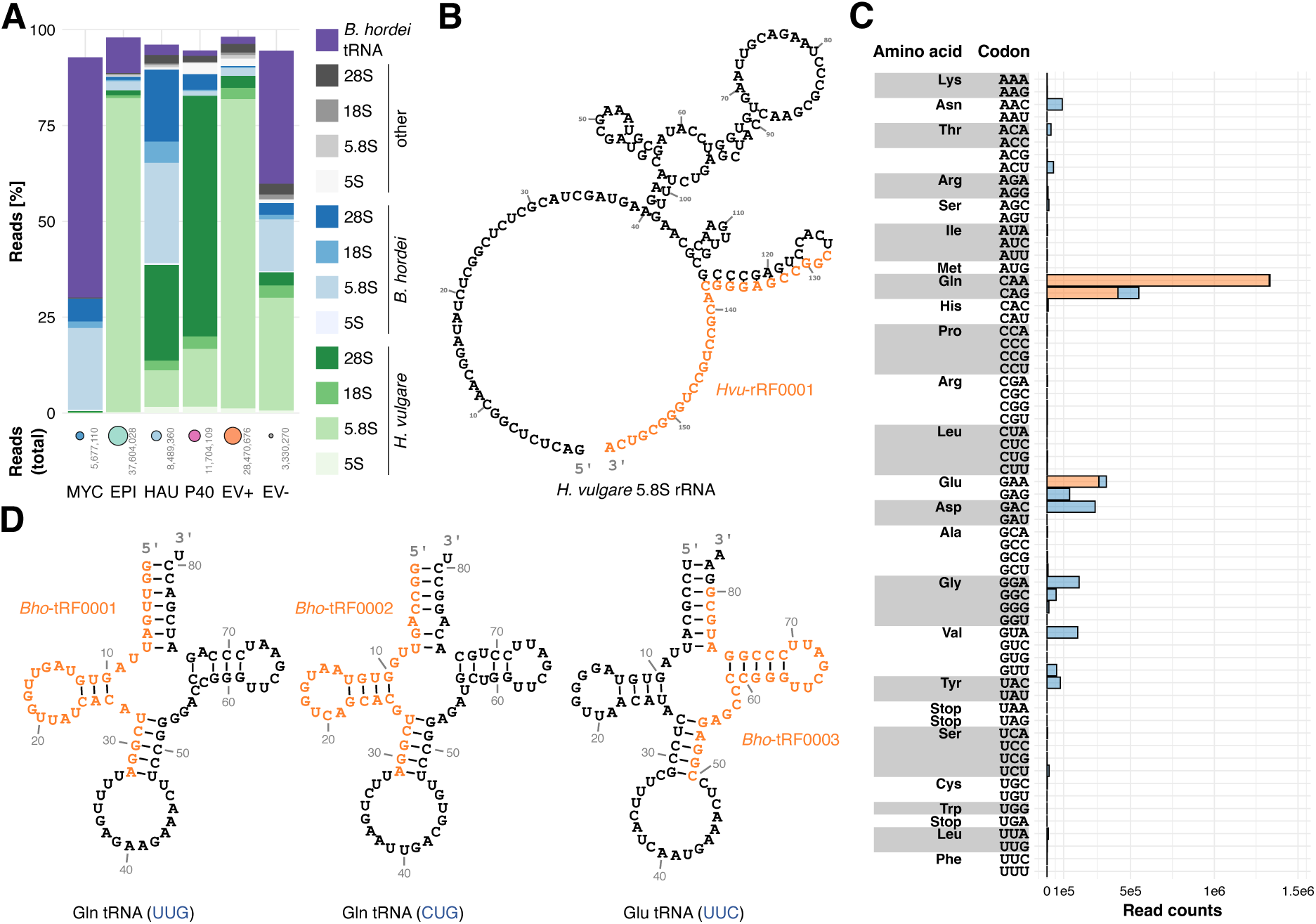
Specific barley 5.8S rRNA- and *B. hordei* tRNA-derived sRNAs are enriched in the 31-33 base reads. We aligned sRNA-seq reads of 31-33 bases in length to the RFAM database using MMSeqs2 (88). (**A**) The stacked bar graph shows the percentage of reads identified as 5S, 5.8S, 18S (small subunit, SSU), 28S (large subunit, LSU), or tRNA, as indicated in the color-coded legend. Green, reads identified as derived from *H. vulgare;* blue, reads identified as derived from *B. hordei* DH14; grey, reads not assigned to either *H. vulgare* or *B. hordei;* purple, reads identified as *B. hordei* tRNA-derived. Epiphytic fungal mycelium (MYC), infected epidermis without mycelium (EPI), fungal haustoria (HAU), microsomes of the epidermis without haustoria (P40), apoplastic extracellular vesicles (EV+), and apoplastic extracellular vesicles of non-infected control plants (EV-). The total number of reads assigned to each sample is given below the bar graph (visualized by circle size). (**B**) Predicted secondary structure of the barley 5.8S rRNA (RFAM accession CAJW010993076.1:c203-48; RNA central accession URS0000C3A4AE_112509), calculated by R2DT in RNA central (https://rnacentral.org) and visualized with Forna (79). The RNA sequence in orange indicates the over-represented 3’ end in the reads from the EPI and EV+ samples. (**C**) Histogram showing the number of reads (Read counts, x-axis) accounting for the *B. hordei* tRNA-derived reads in the sample MYC. The coding amino acid and respective mRNA codons are indicated on the left. The orange portion of the histogram bars indicates the fraction of reads coming from the three most abundant tRNA fragments. (**D**) The three most abundant tRNAs represented in the MYC sample are shown; left, Gln tRNA with UUG anticodon; middle, Gln tRNA with CUG anticodon; right, Glu tRNA with UUC anticodon. The orange-labelled sequences indicate the abundant tRNA fragments.

**Table 1.**
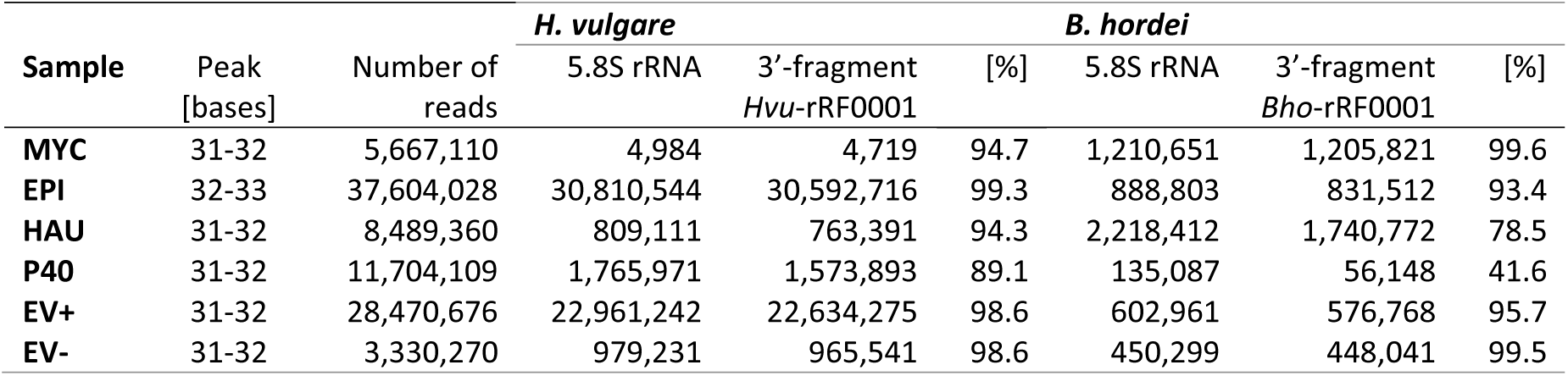
Barley and *B. hordei* sRNAs (rRFs) derived from the 3′-end of the respective 5.8S rRNA.

| Sample | Peak [bases] | Number of reads | *H. vulgare* |  |  | *B. hordei* |  |  |
| --- | --- | --- | --- | --- | --- | --- | --- | --- |
|  |  |  | 5.8S rRNA | 3'-fragment<br>*Hvu*-rRF0001 | [%] | 5.8S rRNA | 3'-fragment<br>*Bho*-rRF0001 | [%] |
| MYC | 31-32 | 5,667,110 | 4,984 | 4,719 | 94.7 | 1,210,651 | 1,205,821 | 99.6 |
| EPI | 32-33 | 37,604,028 | 30,810,544 | 30,592,716 | 99.3 | 888,803 | 831,512 | 93.4 |
| HAU | 31-32 | 8,489,360 | 809,111 | 763,391 | 94.3 | 2,218,412 | 1,740,772 | 78.5 |
| P40 | 31-32 | 11,704,109 | 1,765,971 | 1,573,893 | 89.1 | 135,087 | 56,148 | 41.6 |
| EV+ | 31-32 | 28,470,676 | 22,961,242 | 22,634,275 | 98.6 | 602,961 | 576,768 | 95.7 |
| EV- | 31-32 | 3,330,270 | 979,231 | 965,541 | 98.6 | 450,299 | 448,041 | 99.5 |

**Table 2.**
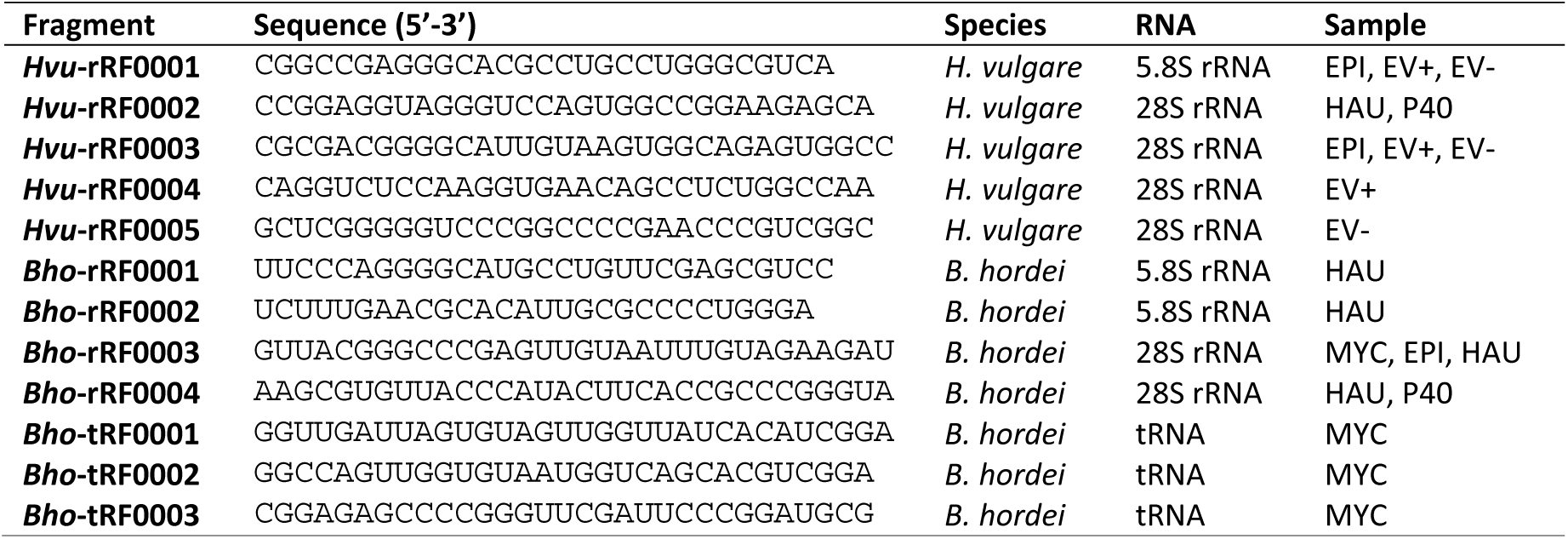
Characteristic *H. vulgare* and *B. hordei* RNA fragments derived from tRNA (tRF) and rRNA (rRF).

In the P40 and HAU samples, which had a broad peak from 27 to 32 bases (Supplementary Figure 1), a large portion of reads was identified as 28S (large subunit, LSU) rRNA-derived sRNAs (Figure 2 and Supplementary Figure 3). P40 had >60% of barley 28S rRNA-originating reads, while HAU reads could be mapped to >25% barley and >20%*B. hordei* 28S rRNA sequences. We aligned these reads in a targeted manner to the 28S LSU rRNAs of *H. vulgare* and *B. hordei* and noted distinct prominent peaks, suggesting the enrichment of specific 28S rRFs in *B. hordei*-infected barley plants (Supplementary Figure 4 and 5, Supplementary Table 5 and 6). In case of the barley 28S rRNA, the first peak was approximately at position 2,100-2,150 (*Hvu*-rRF0002) and the second peak around position 3,760-3,800 (*Hvu*-rRF0003); both peaks were distinctive in the HAU and P40 samples. The peak corresponding to *Hvu*-rRF0002 was shifted to 2,230-2,270 in EV+ samples (*Hvu*-rRF0004), while the peak at 2,100-2,150 was somewhat split and shifted to positions 2,340-2,380 in the EV-sample (*Hvu*-rRF0005). For *B. hordei*, the 28S rRNA-derived sRNAs mapped to a peak at position 430-470 (*Bho*-rRF0003), which is distinctive in the EPI, HAU, and MYC samples. In addition, a second peak at position 1,620-1,660 (*Bho*-rRF0004) appeared in the samples HAU and P40. Notably, we detected a peak at position 2,150-2,180 in the EV+ sample, which is probably due to the high identity between the two 28S rDNA sequences between *H. vulgare* and *B. hordei* in this position (Supplementary Figure 5D).

More than 60% (3,557,695) of the reads from the MYC sample were identified as *B. hordei* tRNA-derived sRNAs (tRNA fragments (tRFs); Figure 2). Intriguingly, >48% of these appear to originate from *B. hordei* tRNAs with anticodons for glutamine (Gln) (Figure 2D). The 5’ moiety of the UUG anticodon Gln tRNA accounted for 1,332,676 reads (37%), 424,142 reads (11.9%) were from the 5’ end and 124,372 (3.5%) from the 3’ end of the CUG anticodon Gln tRNA, and 314,017 reads (8.8%) were identified as the 3’ moiety of the UUC anticodon Glu tRNA. Next, we predicted the putative secondary structures and minimum free energy of the rRNA and tRNA fragments using the Vienna RNAfold webserver (30) to assess if they would be thermodynamically stable (Supplementary Table 7). Notably, *Hvu*-rRF0001 was predicted to form a secondary structure that was particularly stable, forming two double-stranded helices at a free energy of −11.9 kcal mol^−1^ (Supplementary Figure 6). This was markedly more stable than any other theoretical fragment that could be derived from the barley rRNA (between −1.5 and −4.7 kcal mol^−1^) and also than the *B. hordei* 5.8S rRNA-derived fragments (−5.3 and −3.4 kcal mol^−1^, respectively). However, not all abundant rRNA fragments exhibited low free energy (range between −2.4 and −11.9 kcal mol^−1^; Supplementary Table 7). Similarly, the predicted secondary structures of the *B. hordei* tRNA fragments ranged from −1.7 to −10.4 kcal mol^−1^ free energy, overall suggesting that the tRNA and rRNA fragments we observed (Table 2) may assume secondary structures that are thermodynamically stable, but calculated free energy is insufficient evidence to explain the high abundance of most fragments in our dataset.

To assess if specific rRNA fragments occur frequently in infected plants, we then mined publicly available sRNA sequencing datasets from barley, wheat (*Triticum aestivum*), soybean (*Glycine max*), and *Arabidopsis thaliana* under various biotic and abiotic stresses (see Supplementary Table 8 for all accessions). These data showed that read length distributions varied between experiments even within the same species (Supplementary Figure 7). Notably, we did not find striking rRNA enrichment in response to infection or abiotic stress in this subset of sequencing data (Supplementary Figure 8). The fact that another *B. hordei* sRNA-seq dataset (24) also lacks evidence for abundant rRNA suggests that these fragments either appear late in infection (three to four days post inoculation in our samples versus 48 hours post inoculation in (24)) or are only detectable in sampling material enriched with infected host cells. Altogether we identified distinctive barley 5.8S, 28S (both barley and *B. hordei*), and *B. hordei* tRNA-derived sRNAs enriched in *B. hordei*-infected barley samples (Table 2).

### Identification of microRNA-like (milRNA) genes in *B. hordei* and in barley

Next, we used the ShortStack pipeline (Materials and Methods) to annotate and quantify our sRNA samples. Because *bona fide* miRNAs are difficult to predict in fungi (6), we identified microRNA-like RNAs (milRNAs) that satisfy all conditions of miRNAs except detection of the precursor strand. Cumulatively, we identified 2,711 unique *B. hordei* and 35,835 unique barley milRNA loci with this pipeline, accounting for 2,558 and 29,987 unique milRNA sequences, respectively (Supplementary Table 9; Supplementary Files 2-5). Of these, three *B. hordei* milRNAs and 59 barley sequences were classified as *bona fide* miRNAs by ShortStack. For *B. hordei*, most milRNAs were detected in the EPI, MYC, HAU, and EV+ samples (more than 200 milRNAs each), reflecting the abundance of fungal reads in these samples. In case of *H. vulgare*, EPI and MYC samples represented the largest numbers of milRNAs (more than 5,000 each), followed by the EV-, HAU, and P40 samples.

Then, we determined read counts for milRNA loci in both *H. vulgare* and *B. hordei* using the read mapping information from ShortStack (Supplementary Table 10 and 11). We analysed sample relatedness by using non-metric multi-dimensional scaling (NMDS), which collapses multidimensional information into few dimensions (two in this case; Figure 3). In case of *H. vulgare*, the three replicates of MYC, EPI, HAU, and P40 each formed distinct clusters, while the EV+ samples were distinct but only broadly clustered, suggesting stronger variation between replicates. EV-samples did not show clear clustering, as two data points were similar to the EPI and MYC samples, and one was similar to the EV+ samples (Figure 3A). The replicates distributed similarly in case of the *B. hordei* samples, but P40 samples showed a broader distribution and some overlap with the EV+ sample (Figure 3B). For both *H. vulgare* and *B. hordei*, the samples showed comparable Pearson-based hierarchical clustering (Figure 3C and 3D) and clustering trends in principal component analysis (PCA; Supplementary Figure 9); ANOSIM testing confirmed significant sample differentiation in both *H. vulgare* and *B. hordei* (*p* < 1e-4). EV-replicate three, which clustered with EV+ samples, also showed the distinct peak of 31-32-base long reads, enrichment of barley 5.8S rRNAs and *B. hordei* tRNAs, and of 28S rRNA-derived fragments (Supplementary Figure 10). The 31-32-base peak was also visible in EV-replicate two and consisted of >25%*B. hordei* 5.8S rRNA-derived and >50% tRNA-derived sRNAs. This suggests that EV-replicates two and three represent vesicles from unintentionally infected rather than non-infected barley leaves, and that the *H. vulgare* rRFs only occur due to infection with *B. hordei* as the only replicate without apparent *B. hordei* sRNAs also lacks the peak at 31-32 bases. Together with the detection of *B. hordei* rRNA-derived reads of 31-33 bases in length (Figure 2), these analyses suggest that at least one EV-sample was similar to the EV+ samples, another contained a significant number of *B. hordei-derived* sRNAs, and that EV samples generally exhibited high variation hinting at a possible contamination of one or two EV-samples and/or a low signal-to-noise ratio in the EV fractions.

**Figure 3.**
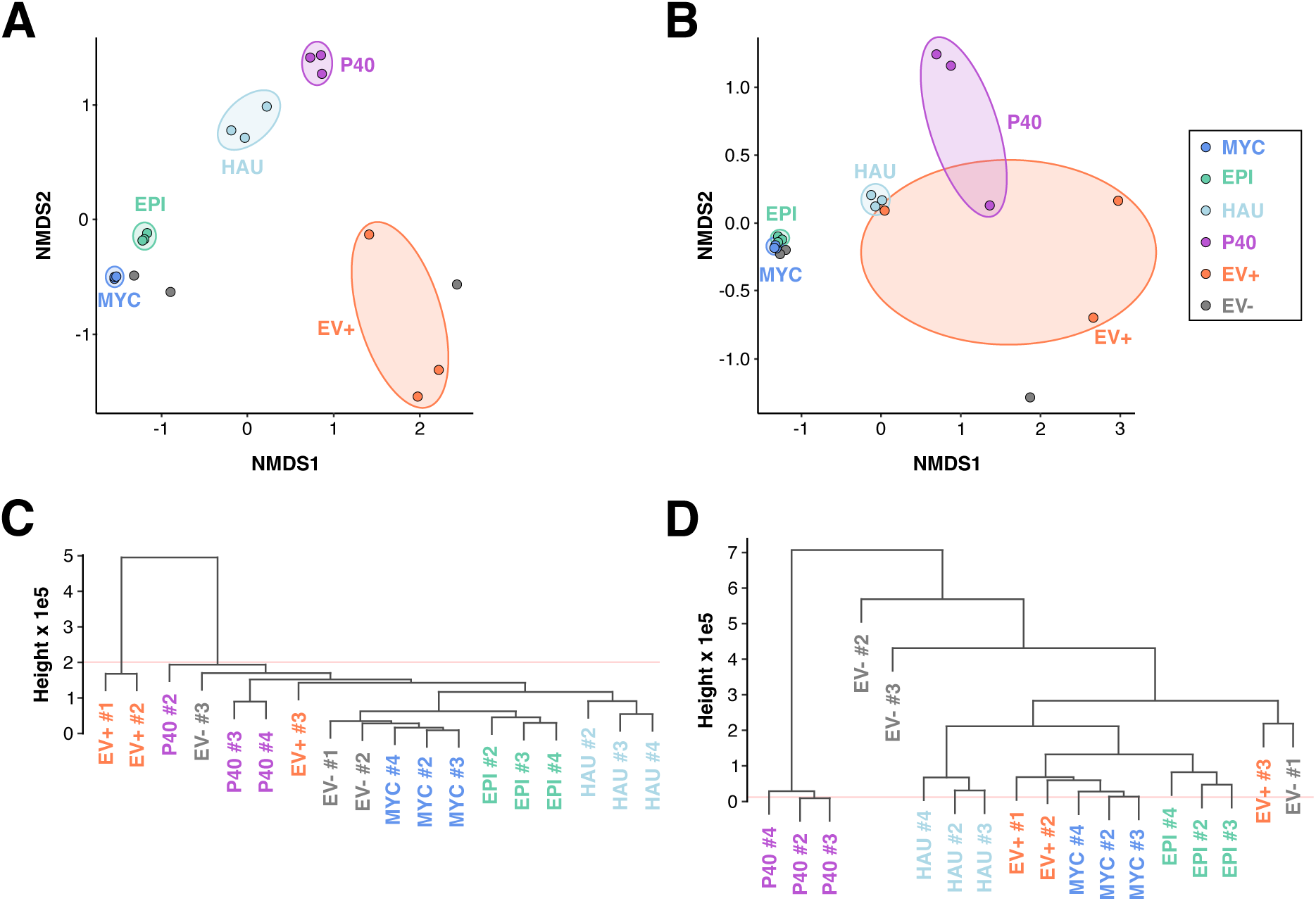
The milRNA content of microsomal samples differs from mycelial, epidermal, and haustorial samples. (**A**) and (**B**). We used non-metric multi-dimensional scaling (NMDS), which collapses multidimensional information into two dimensions to visualize sample similarity. Each data point represents the collapsed milRNA expression data from *H. vulgare* (**A**) and *B. hordei* (**B**). Blue, epiphytic fungal mycelium (MYC); green, infected epidermis without mycelium (EPI); light blue, fungal haustoria (HAU); purple, microsomes of the epidermis without haustoria (P40); orange, apoplastic extracellular vesicles (EV+); grey, apoplastic extracellular vesicles of non-infected control plants (EV-). (**C**) and (**D**) milRNA sample distances based on a Pearson correlation matrix from the milRNA expression data. The pair-wise Pearson correlations were used to calculate a Euclidean distance tree with all samples for *H. vulgare* (**C**) and *B. hordei* (**D**).

### Site-specific accumulation of milRNAs

We performed “weighted gene co-expression network analysis” (WGCNA) to identify milRNAs associated with MYC, EPI, HAU, P40, and EV+ samples for 22,415 *H. vulgare* and 2,711 *B. hordei* milRNAs (Supplementary Figure 11), and then assigned 11,334 *H. vulgare* and 2,325 *B. hordei* milRNAs as enriched in compartments with at least 2.5-fold abundance above the average. 2,496 *H. vulgare* milRNAs were enriched in MYC, 1,137 in EPI, 422 in HAU, 410 in P40, and 615 in EV+ (Figure 4A). In case of *B. hordei*, 519 milRNAs were enriched in MYC, 260 in EPI, 108 in HAU, 59 in P40, and 26 in EV+ (Figure 4B). The samples MYC and EPI also showed considerable overlap of equally abundant milRNAs, as 1,009 *B. hordei* milRNAs and 3,628 *Hvu* milRNAs were more frequent in both samples. Further, 137 *B. hordei* milRNAs were associated with MYC, EPI, and HAU. We used psRNAtarget (31) to predict RNAi targets of *H. vulgare* and *B. hordei* milRNAs. Overall, 3,693 *H. vulgare* milRNAs had putative targets, i.e., 10,488 endogenous and 1,309 *B. hordei* unique transcripts (Figure 4). Ninety (6.8%) of the possible *B. hordei* target genes code for proteins with a predicted secretion peptide. In case of *B. hordei* milRNAs, we found that 1,205 milRNAs could target 581 endogenous genes, 50 (8.6%) of which encode proteins with a predicted secretion peptide, and 1,677 may target transcripts in the host *H. vulgare*. Notably, HAU-specific *B. hordei* milRNAs had 76 potential cross-kingdom targets in *H. vulgare*, but only 23 endogenous targets. These milRNAs exhibited higher average abundance (837 transcripts per million (TPM) on average) than, e.g., MYC-specific milRNAs (59 TPM average) or all milRNAs (181 TPM average). The *B. hordei* milRNAs enriched in HAU and P40 (23 milRNAs, 5,768 TPM on average) exhibited a similar pattern, predicted to target altogether 12 *H. vulgare* genes and only one *B. hordei* gene.

**Figure 4.**
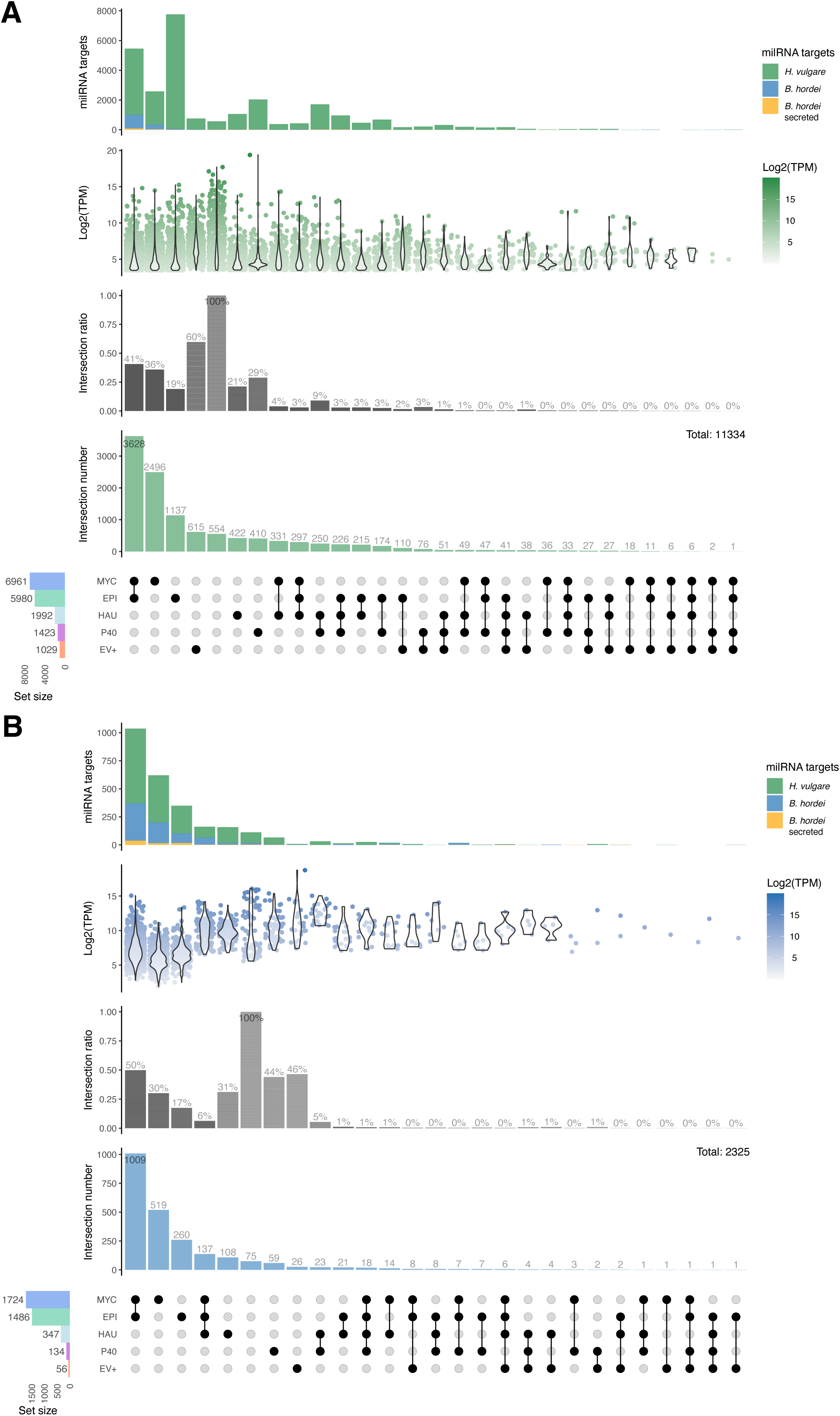
Large sets of milRNAs exhibit site-specific distribution. We calculated site-specific abundance in MYC, EPI, HAU, P40, and EV+ samples for (**A**) *H. vulgare* and (**B**) *B. hordei* using WGCNA and assigned milRNAs to samples exhibiting >2.5-fold enrichment over the average. The bottom panels indicate the set sizes of milRNAs in the respective samples (bar graph, left); blue, epiphytic fungal mycelium (MYC); green, infected epidermis without mycelium (EPI); light blue, fungal haustoria (HAU); purple, microsomes of the epidermis without haustoria (P40); orange, apoplastic extracellular vesicles (EV+). The dots indicate the samples contributing to the respective interaction sets. The second panels from the bottom are bar graphs of the intersection numbers, which are the numbers of milRNAs in the exclusive intersections. The third panels from the bottom show the normalized abundance of the milRNAs in the respective interaction sets as violin plot; the maximum TPM value across samples was used for each data point. The top panels are stacked bar charts of the sum of predicted milRNA target genes in *B. hordei* (blue) and *H. vulgare* (green); *B. hordei* genes encoding a secreted protein are shown in orange.

We then performed global and subset-specific gene ontology (GO) enrichment analysis of the putative cross-kingdom milRNA targets (Figure 5; Supplementary Table 12 and Supplementary Table 13). We found significant GO enrichment in three of the *B. hordei* milRNA cross-kingdom target sets (Figure 5A). Protein phosphorylation-related terms (GO:0006468 and GO:0016773) were enriched between 3.6- and 12.9-fold in all three of the milRNA cross-kingdom target sets (*P*_adj_ < 0.05). “ATP binding” was 1.5-fold enriched (*P*_adj_ < 0.0011) in the predicted cross-kingdom targets of MYC- and EPI-specific *B. hordei* milRNAs. Two GO terms relating to protein K63-linked deubiquitination (GO:0070536 and GO:0061578) were 80.9-fold enriched in the putative cross-kingdom targets of *B. hordei* milRNAs enriched in MYC, EPI, and HAU (*P*_adj_ < 0.05). Only three *H. vulgare* milRNA cross-kingdom target sets, containing MYC-specific, EPI-specific, or MYC- and EPI-specific milRNAs, showed significant enrichment of GO terms (Figure 5B). Two genes targeted by MYC- and EPI-specific milRNAs had the GO term “microtubule-severing ATPase activity” (*P*_adj_ < 0.046; 128.9-fold enrichment), while the term “vacuole” was 6.5-fold enriched in the same set (*P*_adj_ < 0.004). In the MYC-specific set, “ADP binding” was 6.2-fold enriched (*P*_adj_ < 0.04) and “nucleoside-triphosphatase activity” 5.1-fold (*P*_adj_ < 0.02); “ADP binding” was also enriched in the EPI-specific set (11-fold, *P_adj_* < 0.009). The only GO terms enriched in endogenous milRNA targets of *B. hordei* were related to protein phosphorylation (Supplementary Figure 11A), exhibiting 4-fold to 11-fold enrichment (*P*_adj_ < 7e-10) in MYC-, EPI-, MYC- and EPI-, and MYC/EPI/HAU-specific sets. Conversely, we found 69 enriched GO terms in the sets of putative endogenous *H. vulgare* target genes Supplementary Figure 11B; Supplementary Table 12 and Supplementary Table 13). Many of these GO terms relate to regulatory, cell cycle, growth, and developmental processes.

**Figure 5.**
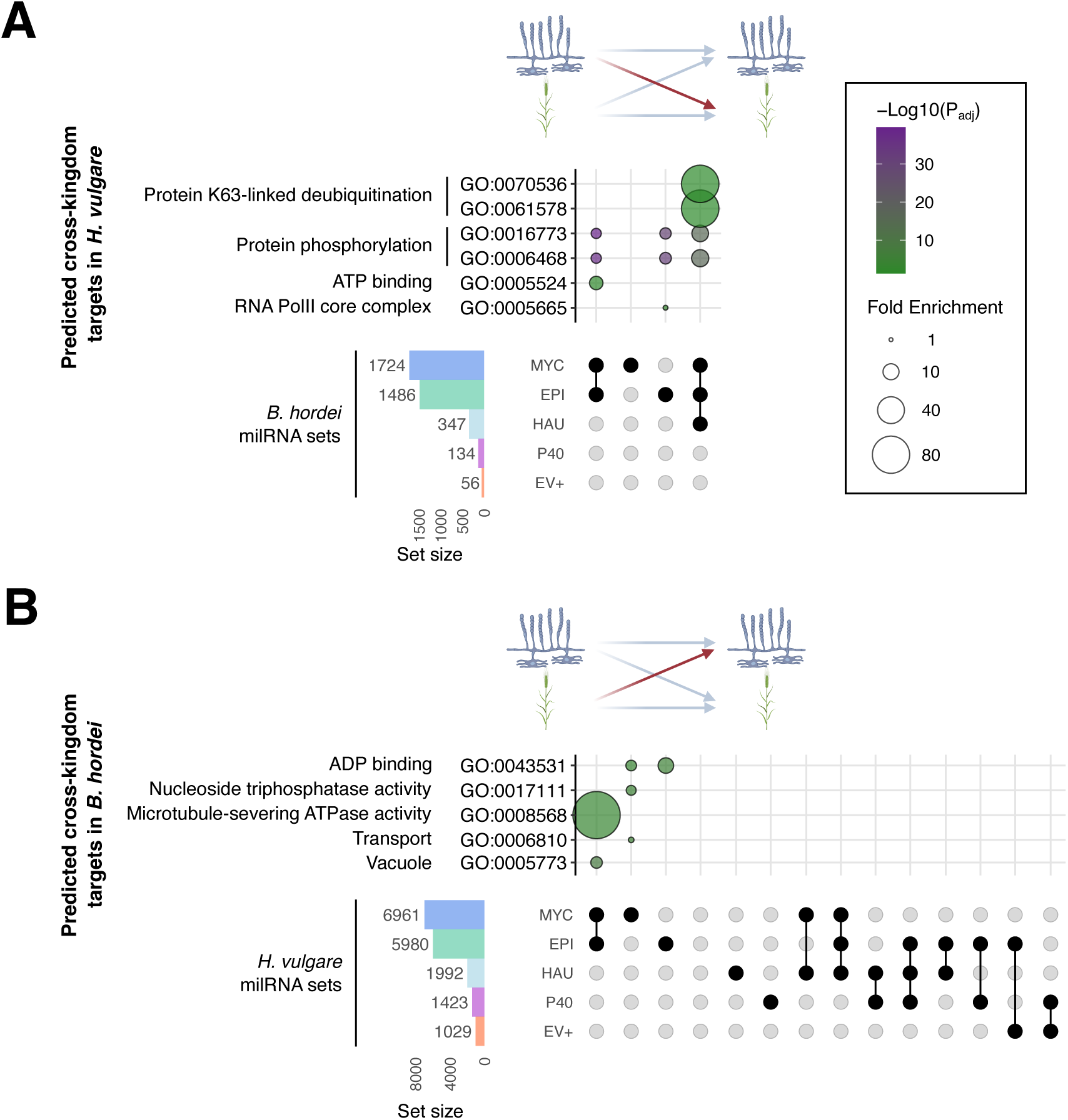
GO enrichment analysis of putative cross-kingdom milRNA targets. We determined all putative targets of the sets of *B. hordei* and *H. vulgare* milRNAs via psRNAtarget (31). We used ShinyGO v0.75 (85) to calculate enriched gene ontology (GO) terms in all milRNA target sets and summarized redundant GO terms with EMBL-EBI QuickGO (https://www.ebi.ac.uk/QuickGO/) on GO version 2022-04-26 and REVIGO (86). (**A**) GO enrichment terms found in putative cross-kingdom targets of *B. hordei* milRNAs in *H. vulgare*. (**B**) GO enrichment terms found in putative cross-kingdom targets of *H. vulgare* milRNAs in *B. hordei*. The GO terms and identifiers are indicated next to the bubble plots. Bubble size indicates fold enrichment of the term in the respective subset, fill color indicates -Log10 of the FDR-adjusted enrichment *P* value. The milRNA subsets are indicated below the bubble plots (see Figure 4 for all subsets). The icons on top of the plot were created with bioRender.com; the blue mycelium indicates *B. hordei* and the green plant barley.

Altogether, we detected that most *B. hordei* and *H. vulgare* milRNAs exhibited site-specific enrichment patterns, suggestive of their infection stage- and tissue-specific induction. Among the putative targets of *B. hordei* milRNAs, genes coding for proteins involved in protein phosphorylation appeared to be consistently overrepresented, both in presumed endogenous and cross-kingdom gene targets. Predicted targets of *H. vulgare* milRNAs showed some enrichment of processes related to microtubule-severing ATPase activity in cross-kingdom *B. hordei* target genes, while endogenously regulating cell cycle, growth, and development.

### *B. hordei* shows signs of phasing in coding genes and retrotransposons

In addition to detecting milRNA candidates, ShortStack (29) can suggest loci containing phased siRNAs (phasiRNAs). Unexpectedly, ShortStack detected that phasiRNAs had occurred in RNAs from 22 *B. hordei* loci in our dataset, present in samples EPI and MYC but none of the other samples (phasing score > 30). Of these, two were found in genes encoding Sgk2 kinases (*BLGH_05411* and *BLGH_03674*), which are abundant in the genome of the fungus (32), including at least one apparent pseudogenized Sgk2 kinase (*BLGH_03674;* Figure 6, Supplementary Figure 12). Another four phasing loci corresponded to coding genes, namely *BLGH_02275, BLGH_00530, BLGH_00532* (encoding proteins of unknown function), and *BLGH_03506* (encoding CYP51/Eburicol 14-alpha-demethylase, involved in ergosterol biosynthesis). The remaining 16 phasing loci were found to be in retrotransposons (Supplementary Table 14). Since ShortStack is not optimized for phasiRNA prediction (29), we used the PHASIS pipeline (33), unitas (34), and PhaseTank (35) to detect evidence of phasing in the genome of *B. hordei* with our sRNA-seq dataset. PhaseTank failed to predict any phasiRNAs with our data. However, we detected 153 putative phasing loci with PHASIS, consisting of 21- or 22-base long phasiRNAs (*P* < 0.0005, Supplementary Table 14). We found that nine phasiRNA loci were located in coding genes of *B. hordei*, of which seven encode Sgk2 kinases. Further, one coding gene subject to phasiRNA enrichment (*BLGH_05762)* encoded a putative secreted protein, and another (*BLGH_00843*) a gene encoding a protein of unknown function. Nineteen phasing loci were in the intergenic space. The majority of phasiRNAs were found in retrotransposons of *B. hordei*, i.e., 42 *Tad*, 7 *HaTad*, 23 *Copia*, 29 *Gypsy*, and 23 *NonLTR* retrotransposons (Supplementary Table 14), accounting for 124 loci altogether. One DNA transposon type *Mariner-2* and one unknown transposon was found as well. The most sensitive detection method with our data was unitas (34), which predicted 1,694 unique phasing loci in the genome of *B. hordei*(Supplementary Table 14). These loci were randomly distributed on the scaffolds of *B. hordei* (Figure 6A). Of these, 135 were located in coding genes and 1,535 in transposable elements, while 24 loci were intergenic (Figure 6C). Of the coding genes, 67 encoded Sgk2 kinases, while seven encoded predicted secreted effector proteins. GO terms related to protein phosphorylation and organonitrogen metabolism were enriched in this set of genes (Figure 6D). The abundance of phasiRNAs in the different sites varied greatly; for example, the reads mapping to the Sgk2 kinase locus *BLGH_03674* were abundant in the MYC and EPI samples, while much less numerous in HAU and P40. Surprisingly, they were also very abundant in the EV+ fraction (Supplementary Figure 12).

**Figure 6.**
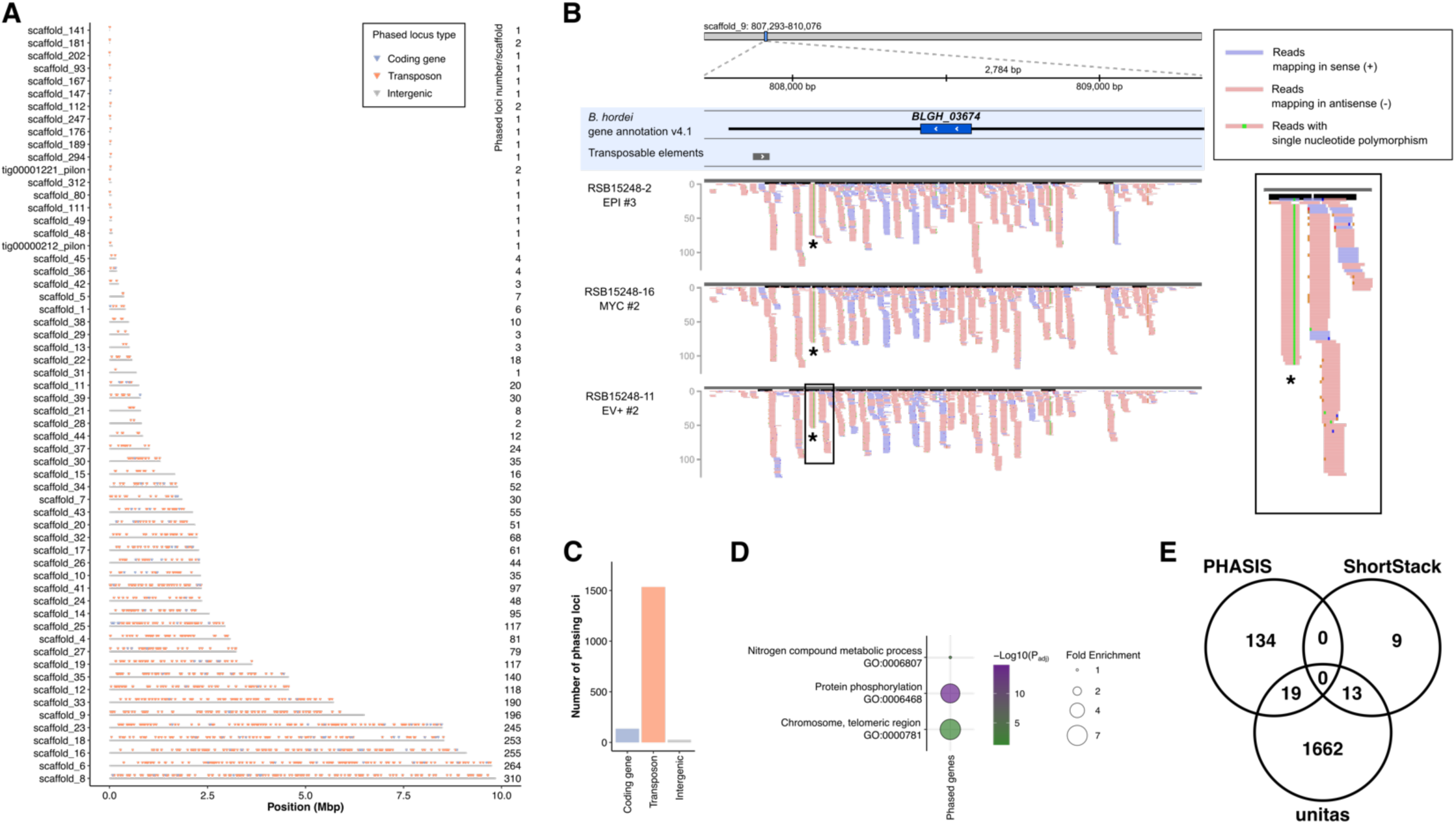
Transposable elements and genes encoding Sgk2 kinases are subject to phasing in *B. hordei*. We identified phasiRNA-rich loci indicative of phasing in the genome of *B. hordei* with ShortStack pipeline v3.8.5-1 (29), the PHASIS pipeline (33), and unitas v1.7.0 (34). (**A**) Global distribution of predicted phasing loci in the genome of *B. hordei* DH14 (28). The x-axis shows the genome position in mega base pairs (Mbp) and the scaffolds are indicated on the y-axis. Triangles denote loci in which phasiRNAs were found with unitas. Blue triangles, phasing loci coinciding with annotated coding genes; orange triangles, phasing loci coinciding with transposable elements; grey triangles, intergenic phasing loci. (**B**) Example of one phased locus in *B. hordei*, containing the gene *BLGH_03674*, the partial open reading frame of a gene encoding a Sgk2 kinase. A subset of representative samples from sites where phasing in this locus was detected is shown (EPI, MYC, and EV+), the full set of samples is displayed in Supplementary Figure 12. From top to bottom, the position on the scaffold, scaffold name and window, gene and transposable element annotations, and sample names are indicated. Red lines indicate reads mapping in sense orientation of the displayed sequence window; blue lines show reads mapping in antisense orientation. A zoom-in is shown on the right. The putative trans-activating RNA (tasiRNA) is indicated with an asterisk. The figure was generated after inspection with Integrative Genome Viewer (IGV; (87)). (**C**) Bar graph summarizing the number of phasing loci types. The x-axis indicates the locus type, i.e., coding gene, transposon, or intergenic; the y-axis shows the number of loci. (**D**) GO enrichment of phased coding genes was calculated with ShinyGO v0.75 (85); terms were summarized to non-redundant GO terms with EMBL-EBI QuickGO (https://www.ebi.ac.uk/QuickGO/) on GO version 2022-04-26 and REVIGO (86). The bubble size indicates fold enrichment of the term in the respective subset, fill color indicates -Log10 of the FDR-adjusted enrichment *P* value. (**E**) A Venn diagram summarizing the overlap of discovered phasing loci with the three methods.

## Discussion

In this work, we sought to characterise the spectrum of sRNAs present in the context of a compatible barley-powdery mildew interaction by isolating and sequencing sRNAs from different “sites” of infected leaves: the epiphytic fungal structures that include the mycelium, runner hyphae, conidiophores and conidia (MYC), the infected barley epidermis including plant tissue and the intracellular fungal haustoria (EPI), a fraction enriched of fungal haustoria (HAU), microsomes obtained after lysis of the epidermal cells (P40), and extracellular vesicles from the apoplast of non-infected (EV-) and infected (EV+) leaves. The first aim was to determine whether the sRNAs differed between the sites. First, we discovered abundant and highly specific fragments derived from 5.8S and 28S rRNA of barley and *B. hordei*, and of tRNAs from *B. hordei* (rRFs and tRFs, respectively; Figure 2 and Table 2). In most cases, these fragments were of discrete lengths (31-33 bases), derived from the 3’ end of the 5.8S rRNA, specific locations within the 28S rRNA, or represented tRNA halves. The fragments exhibited highly specific and prominent enrichment at specific sites of the interaction; thus, they are unlikely to be the product of random RNA exonuclease degradation. Noncoding RNAs including tRNAs and rRNAs preferentially give rise to terminal fragments in animals (36, 37), fungi (38), and plants (39). rRFs have been shown to be involved in ageing in *Drosophila melanogaster* (40), rRNA degradation in response to erroneous rRNAs and UV irradiation stress in *Caenorhabditis elegans* (41, 42), and the response to pathogen infection in black pepper (*Piper nigrum;* (39). Many barley rRFs were abundant in EV+ while not enriched in some EV-samples derived from noninfected plants (Supplementary Figure 10), suggesting their biogenesis is induced by fungal infection in barley. By contrast, *B. hordei* 5.8S rRNA-derived rRFs were particularly abundant in isolated haustoria, suggestive of a role related to the intimate interaction with the host plant.

tRFs appear to be involved in multiple regulatory processes in plants (43). They may play a role in translation inhibition, since 5’-terminal oligoguanine-containing tRF halves of *A. thaliana* can inhibit translation *in vitro* (44). They accumulate under various stresses including phosphate starvation in barley and *A. thaliana* (45, 46) and *Fusarium graminearum* infection in wheat (47). Further, tRNA halves and 10-16 base tiny tRFs represent substantial portions of the sRNA pool in *A. thaliana* and may not be random tRNA degradation products (48). We observed fungal rather than plant tRFs of 31-32 bases specifically in isolated mycelium (Table 2). Some 28-35-base long tRFs are abundant in appressoria of the rice blast pathogen *Magnaporthe oryzae* (49), suggesting infection stage- or tissue-specific roles for these fragments in plant-pathogenic fungi. Intriguingly, the nitrogen-fixing bacterium *Bradyrhizobium japonicus* exchanges specific tRFs with its host plant soybean, hijacking the host RNAi machinery and supporting nodulation (50). While our data does not demonstrate *B. hordei* 31-32 base tRF exchange with the host, we detected tRFs in both infected epidermis and one EV-sample (Figure 2; Supplementary Figure 10), which hints at some tRFs occurring in vesicles and infected host cells.

An important question is how such rRNA and tRNA fragments arise and how they are specifically enriched in plants and fungi. Both rRFs and tRFs can be generated by Dicer and loaded onto Ago. For example, 18-26 base long tRFs accumulate in a DCL1-dependent manner and mediate Ago-driven cleavage of retrotransposons in *A. thaliana* (51), and 23-base-long rRFs derived from the 5′ end of 5.8S rRNA appear to be associated with AGO1 in *P. nigrum* (39). In the fungus *Neurospora crassa*, the production of rRFs requires the RNA-dependent RNA polymerase QDE-1, the helicase QDE-3, and DCL (38). However, the fragments we observed were longer than 26 nucleotides. Thus, they are unlikely to be associated with Dicer and Ago proteins. Instead, these specific tRFs and rRFs may have been generated by specific RNases, like the RNase A Angiogenin in humans (52), the *Saccharomyces cerevisiae* RNase T2 Rny1p (53), and plant RNase T2 (54–56, 47). RNase T2 most likely recognizes and hydrolyses single-stranded rRNA/tRNA loops, which could explain why the ends of the tRFs and rRFs we observed are always found in single-stranded loops (Figure 2). However, it is not clear why specific rRFs and tRFs accumulate while the remaining rRNA and tRNA molecules disappear. We found that these fragments potentially form novel secondary structures, which may render them inaccessible to RNA exonucleases, and that the barley 5.8S rRF may be energetically more stable than other fragments derived from the same molecule (Supplementary Figure 6). However, this is not the case for all rRFs and tRFs we found, suggesting that predicted thermodynamic stability alone cannot explain their high abundance. Future research will show if plants and pathogens repurposed specific fragments of these evolutionary ancient rRNA and tRNA molecules specifically towards intra- or cross-species communication.

Next, we turned our attention to miRNAs and milRNAs; in particular we attempted to predict possible targets for endogenous and putative cross-kingdom gene regulation by RNAi. We found that barley milRNAs are predicted to target more endogenous genes than fungal genes (Figure 5), while *B. hordei* milRNAs are predicted to target more genes cross-kingdom in barley. This trend was particularly evident in haustoria (76 of 108 milRNAs had calculated cross-kingdom targets). The most frequent process associated with putative gene targets for *B. hordei* milRNAs in samples derived from epiphytic mycelium and infected epidermis is protein phosphorylation. Protein phosphorylation is important during plant defence, especially for signal transduction (57). In support of this notion, kinases have been reported as sRNA targets before. For example, *B. cinerea* sRNAs target *A. thaliana* MAP kinases and cell wall-associated kinases (11). Similarly, sRNAs from the wheat stripe rust *Puccinia striiformis* f.sp. *tritici* are predicted to target many kinase genes in wheat (58), and *Fusarium oxysporum mil-R1* interferes with kinases of its host, tomato (59). Although fewer, there are also potential cross-kingdom targets for barley milRNAs. For example, barley milRNAs were predicted to target *B. hordei* genes related to microtubule-severing ATPase activity, ADP binding, and nucleoside triphosphatase activity (Figure 5). Microtubules play key roles in tip growth of hyphae in fungi, and the severing ATPase activity is essential for microtubule organization (60, 61). It is therefore possible that plant miRNAs have evolved to interfere with microtubule dynamics in the fungal pathogen.

Remarkably, we also detected *B. hordei* milRNAs in EV+ samples. These milRNAs are unlikely to be contaminations because a considerable number (26 in total) of *B. hordei* milRNAs that were specifically enriched in this sample type (Figure 4B). In principle, these EV+-enriched *B. hordei* milRNAs could originate from *B. hordei-derived* microsomes. However, the apoplastic space is physically separated from the main plant-fungus interaction site (the interface between the extrahaustorial membrane and the plant cell wall) by the haustorial neck band (62), which makes vesicle diffusion of fungal origin into the apoplast unlikely. In principle, broken cells or breaking of cells or the haustorial neck during sample preparation are another possibility for the origin of these EVs. Also, during the vacuum infiltration process to obtain the apoplastic fluid, *B. hordei* EVs could be washed off the leaf surface and the mycelium. However, there was limited co-enrichment of milRNAs in MYC and EV+ samples (Figure 4), and the EV+-specific milRNAs preclude this scenario. Therefore, it is possible that *B. hordei* milRNAs hijack barley EVs in a similar manner as the turnip mosaic virus does in *Nicotiana benthamiana* (63). The virus uses this mechanism to disperse its genetic material. However, this scenario would require the transition of fungal sRNAs through several stages of endo- and exocytosis.

PhasiRNAs have been well-documented in plants where they are thought to be the result of complex processes of controlled degradation of double-stranded RNAs; they play roles in a wide variety of biological functions including development and plant immunity (64). Conversely, there is only very limited information about phasiRNA in fungi: to the best of our knowledge the only case where phasiRNAs have been reported in the fungal kingdom is *Sclerotinia sclerotiorum* infected by a hypovirus (8). We were, therefore, surprised to find a clear signature of phasiRNA mapping to the fungal genome in *B. hordei* using ShortStack (Figure 6). We subsequently succeeded in detecting phasiRNAs using two other independent algorithms (unitas and PHASIS; Figure 6). The different pipelines detected somewhat different sets of phased loci in the *B. hordei* genome, which likely reflects the different mapping protocols. PHASIS exhibited the lowest overlap in phasing locations between the three tools; this is probably because PHASIS does not use already existing mapping files but performs its own mapping procedure instead. In any case, the low degree of overlap highlights the suboptimal performance of these tools for the identification of phasiRNA in filamentous fungi, and improvements may need to be implemented to determine complete phasiRNA complements in these organisms. The vast majority of phasiRNAs mapped to repetitive elements in the *B. hordei* genome, i.e., transposons and the previously described abundant *Sgk2* kinase loci (32). This may result from the involvement of phasiRNAs in controlling the expression of such genomic elements in the absence of the otherwise highly conserved repeat-induced point mutation (RIP) pathway (65, 28). In animals, it is thought that PIWI-associated RNAs (that show analogous phasing to plant phasiRNAs) also function to control transposons in *Drosophila* (66). Nevertheless, according to our data, some protein-coding gene loci in *B. hordei* show the presence of phased sRNAs as well, raising the possibility that the expression of these genes may be subject to regulation by RNAi. The presence of predicted phasiRNAs of fungal origin in EVs of the leaf apoplast from infected barley leaves (EV+) is intriguing. It is difficult to rationalise what function, if any, these may have in modulating plant gene expression at remote sites. We note, however, that trans-acting small interfering RNA3a RNAs (tasi-RNA) derived from the *A. thaliana TAS3a* locus and synthesized within three hours of pathogen infection may be an early mobile signal in systemic acquired resistance (67). It is thus conceivable that trafficking of sRNAs of fungal origin may follow the same route and may target plant gene expression (24), and that fungal phasiRNAs in plant EVs are simply unintended by-products of this track. Overall, our findings further support the notion that phasing exists in the fungal kingdom and additionally provide evidence that transposable elements in *B. hordei* are subject to sRNA-directed post-transcriptional regulation.

## Material and Methods

### Plant cultivation

Plants of barley (*H. vulgare* cv. Margret) were grown in SoMi513 soil (HAWITA, Vechta, Germany) under a long day cycle (16 h light period at 23 °C, 8 h darkness at 20 °C) at 60-65% relative humidity and 105-120 μmol s^−1^ m^−2^ light intensity. Seven-day-old barley plants were inoculated with *B. hordei* strain K1; the plants were transferred to growth chambers with a long day cycle (12 h light at 20 °C, 12 h dark at 19 °C) at ca. 60% relative humidity and 100 μmol s^−1^ m^−2^.

### Isolation of small RNAs

Seven-day-old barley was inoculated with *B. hordei* strain K1. At four days after inoculation, the primary leaves were harvested to isolate mycelia, epiphytic, haustoria, and microsomes from infected epidermis (P40). The dissection of the tissues and fractions was carried out as previously described (68). Briefly, this consisted of dipping the excised leaves in cellulose acetate (5% w/v in acetone), letting the acetone evaporate for a few minutes, peeling first the epiphytic structures (MYC; this contained epiphytic mycelia, conidia and spores), then dissecting the adaxial barley epidermis (EPI).

A portion of the epidermis samples were subsequently used to extract haustoria (HAU) and epidermis microsomes (P40). First, the plant cell walls were digested by incubating the epidermis in “Onozuka” R-10 cellulase (YAKULT pharmaceutical - Duchefa Biochemie BV, Haarlem, The Netherlands; dissolved in 2% w/v in potassium vesicle isolation buffer; 20 mM MES, 2 mM CaCl_2_, 0.1 M KCl, pH 6.0; (69)) for 2 h at 28 °C on a rotary shaker (80 rpm). The digested epidermis samples were then filtered through a 40 μm nylon mesh sieve and centrifuged for 20 min at 200 *g*. The pellet containing the haustoria was resuspended in 260 μL potassium vesicle isolation buffer and stored at −80 °C (HAU); ~ 1 to 2 × 10^6^ haustoria were obtained per sample. The supernatant was centrifuged again at 10,000 *g* for 30 min at 4 °C, and the pellet discarded. The resulting supernatant was centrifuged one last time at 40,000 *g* at 4 °C for 1 h; the pellet containing the microsomes was stored at −80 °C (P40). Apoplastic wash fluid was extracted from barley plants at three days after inoculation with *B. hordei* strain K1. Trays of both inoculated and non-inoculated plants were covered with lids and incubated prior to apoplastic wash fluid extraction. Approximately 30 g of leaf fresh weight were collected and vacuum infiltrated with potassium vesicle isolation buffer. Excess buffer was carefully removed and the leaves were placed with the cut ends down in 20 mL syringes. The syringes were inserted into 50 mL centrifuge tubes. Apoplastic wash fluid was collected at 400 *g* for 12 min at 4 °C. Cellular debris was first removed by passing the apoplastic wash fluid through a 0.22 μm syringe filter and further by centrifugation at 10,000 *g* for 30 min at 4 °C. A crude extracellular vesicle fraction was isolated from apoplastic wash fluid based on a previously established protocol (69). The apoplastic wash fluid was centrifuged for 1 h at 40,000 *g* (4 °C) in a swinging bucket rotor to collect extracellular vesicles (EV- and EV+). The extracellular vesicle pellet was resuspended in 50 μL 20 mM Tris-HCl (pH 7.5).

The MYC and EPI samples were ground in liquid nitrogen with quartz sand in a chilled pestle and mortar and RNA was extracted from all samples using TRIzol (Thermo Scientific, Schwerte, Germany) as described by the manufacturer. The RNA from all other samples were extracted by resuspending the frozen samples directly into the TRIzol reagent, and proceeding as described. The quantity and quality of the RNA was measured by spectrophotometry (NanoDrop, Thermo Scientific) and spectrofluorimetry (Qubit, Thermo Scientific).

### sRNA sequencing and data processing

RNA samples were quantified via the Qubit RNA HS Assay (Thermo Scientific), and sized with a High Sensitivity RNA ScreenTape (Agilent, Santa Clara, CA, USA). Library preparation was performed with RealSeq-Dual as recommended by the manufacturer (RealSeq Biosciences, Santa Cruz, CA, USA; (70)) with 100 ng of RNA for each sample. Half the volume of each library was amplified by 20 cycles of polymerase chain reaction. Libraries from all samples were pooled for sequencing in the same flow cell of a NextSeq single-end 75 nt reads run. FastQ files were trimmed of adapter sequences by using Cutadapt (71) with the following parameters: cutadapt -u 1 -a TGGAATTCTCGGGTGCCAAGG -m 15. We further performed quality trimming of reads with Trimmomatic v0.39 (72). FASTQ read data were inspected using FastQC v0.11.5 (Babraham Bioinformatics, Cambridge, UK). FASTQ/FASTA files were parsed with SeqKit v2.1.0 (73).

### Read length distribution analysis

We determined read length counts with custom BASH scripts and plotted these data using ggplot2 v3.3.4 (Wickham 2009) in R v4.1.2 (R Core Team 2021). Reads were mapped to the reference genomes of *H. vulgare* (Hordeum_vulgare.IBSC_v2; (27)) and *B. hordei* DH14 v4 (28) using bowtie within the ShortStack pipeline v3.8.5-1 (29). We used SAMtools v1.9 (74) in conjunction with custom BASH scripts to parse read mapping statistics. Read counts by genomic features, i.e., coding genes, transposable elements, and milRNA genes identified by ShortStack, were determined using featureCounts v2.0.1 (76). *H. vulgare* transposable elements were identified with RepeatMasker v4.0.7 (http://www.repeatmasker.org) using the repeat database version RepBase-20170127 and ‘-species Hordeum’. The data were plotted with ggplot2 v3.3.4 (77) in R v4.1.2 (78).

### Read BLAST searches

We used a custom python script to generate FASTA files containing reads of 21-22 bases, 31-32 bases, and 32-33 bases, respectively, and then deduplicated reads using clumpify.sh of the BBmap package (https://jgi.doe.gov/data-and-tools/software-tools/bbtools/). We downloaded the ribosomal and transfer RNA molecules deposited in the RFAM database (http://ftp.ebi.ac.uk/pub/databases/Rfam/CURRENT/fasta_files/) in October 2021. All reads were aligned to the RFAM databases using MMSeqs2 v9.d36de (Steinegger and Söding 2017). Reads aligning to RF02543 (28S rRNA) were subsequently aligned to the 28S rRNA sequences of *H. vulgare* (RNAcentral accession URS0002132C2A_4513) and *B. hordei* (URS0002174482_62688), respectively. Read alignment coverage was determined via BEDtools v2.25.0 (75). Bar graphs and histograms displaying these data were plotted with ggplot2 v3.3.4 (77) in R v4.1.2 (78).

### Structure and free energy analysis of rRNA and tRNA fragments

We obtained rRNA and tRNA structures predicted by R2DT from RNA central (https://rnacentral.org) and visualized the structures with Forna (79). We further used Vienna RNAfold v2.4.18 webserver (30) to calculate secondary structures and their minimum free energy (MFE) of 5.8S rRNAs, rRNA, and tRNA fragments.

### Identification of milRNAs

We used ShortStack pipeline v3.8.5-1 (29) to predict milRNAs in the genomes of *H. vulgare* (Hordeum_vulgare.IBSC_v2; (27)) and *B. hordei* DH14 (28). Small RNAs with a Dicer Call cut-off of N15 were considered as microRNA-like (milRNA) RNAs. We collected non-redundant *B. hordei* and *H. vulgare* miRNAs and milRNAs in GFF and FASTA formats in Supplementary files 2-5. featureCounts v2.0.1 (76) was used to determine milRNA locus-specific read counts.

### Quantification and clustering of milRNA expression

We first analysed similarities/differences between samples using several statistical approaches. Non-metric multi-dimensional scaling (NMDS) collapses multidimensional information into fewer dimensions and does not require normal distribution. The stress value represented the statistical fit of the model for the data. Stress values equal to or below 0.05 indicated a good model fit. We complemented the NMDS analysis with metric multi-dimensional scaling (MDS), principal component (PC) analysis, and Pearson coefficient correlation (PCC)-based hierarchical clustering. In addition, non-parametric rank-based ANOSIM (analysis of similarity) tests were performed to determine statistical differences between samples.

Weighted gene coexpression network analysis (WGCNA; (80)) was performed using read counts mapping to milRNA loci. First, we filtered out the non-expressed genes (cut-off TPM < 1), leaving 22,415 *H. vulgare* and 2,217 *B. hordei* milRNAs for the construction of the coexpression networks. The scale-free network distribution required determination of the soft threshold *β* (12 for *H. vulgare* and 16 for *B. hordei*) of the adjacency matrix. A module correlation of 0.1 with either sample was the cut-off for identifying milRNAs enriched in the samples MYC, EPI, HAU, P40, and EV+; we then refined assignment of milRNAs to samples at >2.5-fold above average expression across samples. We used pairwise differential expression analysis with DESeq2 (81), EdgeR (82), and limma-VOOM (83) to confirm the overall expression trends of milRNAs from *H. vulgare* and *B. hordei*. Intersections were analyzed by UpSetR plots using ComplexUpset (84).

### Small RNA target prediction and GO enrichment of milRNA target sets

We used psRNAtarget (https://www.zhaolab.org/psRNATarget) Schema V2 2017 release (31) at an expectation value cut-off of 2. We performed GO enrichment on putative milRNA target gene sets using ShinyGO v0.75 (85); *P* values indicated were calculated via false discovery rate (FDR) correction (*P*_adj_). GO terms were summarized by removing redundant GO terms with EMBL-EBI QuickGO (https://www.ebi.ac.uk/QuickGO/) on GO version 2022-04-26 and REVIGO (86) with Gene Ontology database and UniProt-to-GO mapping database from the EBI GOA project versions from November 2021.

### PhasiRNA and tasiRNA detection in *B. hordei*

We used ShortStack pipeline v3.8.5-1 (29), the PHASIS pipeline (33), unitas v1.7.0 (34), and PhaseTank v1.0 (35) to detect evidence of phasing in the genome of *B. hordei* with our small RNA-seq dataset. Detected phasing sites were concatenated to non-redundant phasing loci using BEDtools v2.25.0 (Quinlan and Hall 2010). Manual inspection of phasing loci was done using Integrative Genomics Viewer (87).

## Abbreviations

Ago: Argonaute
Dcl: Dicer-like
EPI: epidermis (colonized by *B. hordei* but epiphytic fungal mycelium removed)
dsRNA: double-stranded RNA
EV: extracellular vesicle
FDR: false discovery rate
GO: gene ontology
HAU: haustoria of *B. hordei*
milRNA: micro RNA-like
miRNA: micro RNA
MYC: mycelium of *B. hordei*
phasiRNA: phased siRNA
P40: microsomes from *B. hordei*-infected epidermis
RNAi: RNA interference
rDNA: ribosomal DNA
rRF: ribosomal RNA-derived small RNA fragment
rRNA: ribosomal RNA
sRNA: small RNA
siRNA: small interfering RNA
tasiRNA: trans-acting short interfering RNA
TPM: transcripts per million
tRF: transfer RNA-derived small RNA fragment
tRNA: transfer RNA

## Author contributions

P.D.S. conceived and conceptualized the project, generated materials (HAU, P40, EPI, MYC), and performed initial analysis of the sRNA-seq data. S.K. contributed to initial data analysis and milRNA identification, performed read length distribution analysis in the samples from this study and published data, analysed milRNA expression data, contributed to GO enrichment analysis, did phasiRNA detection in *B. hordei*, and designed figures. M.S. contributed to milRNA identification, read length distribution analysis, secondary structure predictions of RNA fragments, weighted correlation network analysis of milRNAs, performed milRNA target predictions, and gene ontology enrichment analysis. H.T. generated materials from EV samples. S.K., H.T., and P.D.S. drafted the manuscript, all authors edited the document. P.D.S. and R.P. provided reagents, funds, and laboratory space. All authors read and agreed to the final version of the manuscript.

## Data availability statement

The sRNA sequencing data are deposited at NCBI/ENA/DDBJ under project accession PRJNA809109; the SRA accessions are listed in Supplementary Table 1. The *B. hordei* and *H. vulgare* miRNAs and milRNAs are available in GFF format and FASTA format in Supplementary files 2-5.

## Code availability statement

All computational tools and their versions are described in the respective methods sections, and were run in default mode with modifications indicated. Custom codes for our analysis pipeline are available at https://github.com/stefankusch/smallRNA_seq_analysis.

## Acknowledgments

We thank Blake Meyers for advice on the usage of PHASIS, and members of the DFG-funded FOR5116 consortium for critical feedback. The analysis was performed with computing resources granted by RWTH Aachen University under project ID rwth0146.

## Supplementary Figure legends

**Supplementary Figure 1. Each sample type exhibits a distinctive sRNA read size distribution.** (**A**) We determined the sRNA read counts for read lengths between 15 and 40 bases and plotted the read length (x-axis) against the respective number of reads (y-axis). Three replicates were analyzed for each of our six sample types: Epiphytic fungal mycelium (MYC), infected epidermis without mycelium (EPI), fungal haustoria (HAU), microsomes of the epidermis without haustoria (P40), apoplastic extracellular vesicles (EV+), and apoplastic extracellular vesicles of non-infected control plants (EV-). (**B**) We mapped sRNA sequencing reads of 15 and 40 bases long to the reference genomes of *H. vulgare* IBSCv2 (27) and *B. hordei* DH14 (28). The stacked bar graph shows the read counts (y-axis) for the respective read size (x-axis) from the three replicates. Green, reads mapping to the *H. vulgare* genome; blue, reads mapping to the *B. hordei* DH14 genome; grey, reads that did not map to either of the two genomes.

**Supplementary Figure 2. The majority of reads mapping to *B. hordei* or *H. vulgare* originate from transposable elements or coding genes.** We assigned reads aligning to the genomes of (**A**) *B. hordei* or (**B**) *H. vulgare* to the features mRNA (coding genes; light blue/green), milRNA loci identified in this study (blue/green), and transposable elements (dark blue/green). We plotted the read length (x-axis) against the respective number of reads (y-axis). The six sample types were epiphytic fungal mycelium (MYC), infected epidermis without mycelium (EPI), fungal haustoria (HAU), microsomes of the epidermis without haustoria (P40), apoplastic extracellular vesicles (EV+), and apoplastic extracellular vesicles of non-infected control plants (EV-).

**Supplementary Figure 3. A broad range of reads originates from rRNAs in haustoria and microsomal fractions.** (**A**) We aligned all reads to the 5.8S (red), 18S (blue), and 28S (green) rDNA sequences of *B. hordei* and *H. vulgare*. The stacked bar graph shows the read counts (y-axis) for the respective read size (x-axis) from the three replicates. Three replicates were analyzed for each of our six sample types: Epiphytic fungal mycelium (MYC), infected epidermis without mycelium (EPI), fungal haustoria (HAU), microsomes of the epidermis without haustoria (P40), apoplastic extracellular vesicles (EV+), and apoplastic extracellular vesicles of non-infected control plants (EV-). Colors indicate reads aligning to the different rDNAs: Light blue, *B. hordei* 18S rDNA (RNAcentral accession URS000021D3E6_2867405); dark blue, *H. vulgare* 18S rDNA (URS0000AF30DE_112509); light green, *B. hordei* 28S rDNA (URS0002174482_62688); dark green, *H. vulgare* 28S rDNA (URS000212856A_112509); light red, *B. hordei* 5.8S rDNA (URS00006663F0_546991); dark red, *H. vulgare* 5.8S rDNA (URS0000C3A4AE_112509); grey, reads that did not align with any rDNA sequence. (**B**) Alignment of sRNA sequencing reads of 27-32 bases in length from HAU to the *B. hordei* 5.8S rDNA (154 bases in length). The graph shows the cumulative number of reads from the three replicates (y-axis) mapping to each position of the *B. hordei* 5.8S rDNA (x-axis). (**C**) Secondary structure of the *B. hordei* 5.8S rRNA (RFAM accession CAUH01009408.1:1222-1375; RNA central accession URS00006663F0_546991) predicted by R2DT in RNA central (https://rnacentral.org) and visualized with Forna (79). The RNA sequences in orange indicates the over-represented 3’ end in the reads from the HAU sample.

**Supplementary Figure 4. Specific barley 28S rRNA-derived sRNAs are enriched in the 31-33 base long reads.** (**A**) We aligned sRNA sequencing reads of 31-33 bases in length to the *H. vulgare* 28S rDNA (3,853 bases in length). The graphs display the number of reads identified (y-axis) mapping to each position of the barley 28S rDNA (x-axis). Epiphytic fungal mycelium (MYC), infected epidermis without mycelium (EPI), fungal haustoria (HAU), microsomes of the epidermis without haustoria (P40), apoplastic extracellular vesicles (EV+), and apoplastic extracellular vesicles of non-infected control plants (EV-). (**B**) Secondary structure of the 28S rRNA of *H. vulgare* (RFAM accession CAJW010993076.1:203-48; RNA central accession URS000212856A_112509) predicted by R2DT in RNA central (https://rnacentral.org). The sequence stretches shown in orange indicate the overrepresented 28S rRNA fragments *Hvu*-rRF0002 and *Hvu*-rRF0003 from the HAU and P40 samples. *Hvu*-rRF0003 fragments were also detected in EV+ and EV-samples, in addition to *Hvu*-rRF0004 and *Hvu*-rRF0005. (**C**) Sequences of the enriched 28S rRNA fragments of *H. vulgare* in FASTA format.

**Supplementary Figure 5. Specific *B. hordei* 28S rRNA-derived sRNAs are enriched in the 31-33 bases long reads.** (**A**) We aligned sRNA sequencing reads of 31-33 bases to the *B. hordei* 28S rDNA (3,564 bases). The graphs display the number of reads identified (y-axis) mapping to each position of the *B. hordei* 28S rDNA (x-axis). Epiphytic fungal mycelium (MYC), infected epidermis without mycelium (EPI), fungal haustoria (HAU), microsomes of the epidermis without haustoria (P40), apoplastic extracellular vesicles (EV+), and apoplastic extracellular vesicles of non-infected control plants (EV-). (**B**) Secondary structure of the 28S rRNA of *B. hordei* (RFAM accessions CAUH01009223.1:1667-1 and CAUH01013050.1:1126-1; RNA central accession URS000C6B9A3_546991; full structure was reconstructed from these partial fragments) predicted by R2DT in RNA central (https://rnacentral.org). The RNA stretches shown in orange indicate the over-represented 28S rRNA fragments *Bho*-rRF0003 and *Bho*-rRF0004 from the Myc, Epi, Hau, and P40 samples. The fragment *Hvu*-rRF0004 is indicated in the EV+ plot. (**C**) DNA sequences of the enriched 28S rRNA of *B. hordei* fragments in FASTA format. (**D**) Pairwise alignment between *Hvu*-rRF0004 and the respective orthologous *B. hordei* 28S rDNA sequence. Positions in dark blue indicate identity, positions in light blue mismatches between the two sequences. Numbers above the alignment indicate alignment position, numbers on the left the position in the respective rDNA sequence of *H. vulgare* and *B. hordei*. The sequence covered by *Hvu*-rRF0004 is indicated below the alignment.

**Supplementary Figure 6. Predicted secondary structures of rRNA and tRNA fragments and their minimum free energies.** We used the Vienna RNAfold v2.4.18 webserver (30) to calculate secondary structures and their minimum free energy (MFE) of *H. vulgare* and *B. hordei* 5.8S rRNA (**A**), 28S rRNA (**B**), and tRNA (**C**) fragments. (**D**) The cartoon of the 5.8S rRNA structure indicates the position of the respective fragments; orange fragments indicate rRNA and tRNA fragments detected in our sRNA-seq dataset (Table 2) and blue fragments indicate theoretical fragments of identical or similar length used for comparison. The minimum free energies are indicated for each fragment.

**Supplementary Figure 7. Read length distribution profiles vary between sRNA-seq datasets.** We downloaded publicly available sRNA-seq datasets from the NCBI SRA database at https://www.ncbi.nlm.nih.gov/sra (accessions summarized in Supplementary Table 8) and analyzed the read length profiles of reads between 15 and 40 bases in length. The x-axis shows number of reads and the y-axis the read length in bases. We examined datasets from the following samples: (**A**) *H. vulgare* infected with *B. hordei* at 0, 24, and 48 hpi (24); (**B**) *H. vulgare* under salt stress (89) and aluminium stress (90), respectively; (**C**) *Triticum aestivum* (wheat) after infection with *B. graminis* f.sp. *tritici* at 12 hpi and under 40 °C heat stress (91); (**D**) *T. aestivum* infected with *Zymoseptoria tritici* at 12 dpi (1); (**E**) *T. aestivum* under 37 °C heat stress, continuous light stress, or UV treatment stress (92); (**F**) *Glycine max* (soybean) during nodulation with the bacterial species *Bradyrhizobium japonicum* at 10 and 20 days after inoculation (50); (**G**) *Arabidopsis thaliana* and *Phaseolus vulgaris* (common bean) during infection with *Sclerotinia sclerotiorum* (93); (**H**) *A. thaliana* after infection with *Verticillium dahliae* and the *V. dahliae* mutant *aly1 aly2* (94); (**I**) *A. thaliana* infected with *Hyaloperonospora arabidopsidis* at 3, 4, and 7 dpi (95); (**J**) *Botrytis cinerea* cultivated *in vitro* (11); (**K**) *A. thaliana* infected with *B. cinerea* at 24, 48, and 72 hpi (11).

**Supplementary Figure 8. Most sRNA-seq datasets do not exhibit read length-specific enrichment of rRNA fragments.** We downloaded publicly available sRNA-seq datasets from the NCBI SRA database at https://www.ncbi.nlm.nih.gov/sra (accessions summarized in Supplementary Table 8) and aligned all reads against the respective 5.8S, 18S, and 28S rDNA sequences of each species. The stacked bar graph shows the read counts (y-axis) for the respective read size (x-axis) from all available replicates. Colors indicate reads aligning to the different rDNAs: Light blue, plant 18S rDNA; dark blue, fungal 18S rDNA or bacterial 16S rDNA; light green, plant 28S rDNA; dark green, fungal 28S rDNA or bacterial 23 rDNA; light red, plant 5.8S rDNA; dark red, fungal 5.8S rDNA or bacterial 5S rDNA; grey, reads that did not align with any rDNA sequence. We examined datasets from the following samples: (**A**) *H. vulgare* infected with *B. hordei* at 0, 24, and 48 hpi (24); (**B**) *H. vulgare* under salt stress (89) and aluminium stress (90), respectively; (**C**) *Triticum aestivum* (wheat) after infection with *B. graminis* f.sp. *tritici* at 12 hpi and under 40 °C heat stress (91); (**D**) *T. aestivum* infected with *Zymoseptoria tritici* at 12 dpi (1); (**E**) *T. aestivum* under 37 °C heat stress, continuous light stress, or UV treatment stress (92); (**F**) *Glycine max* (soybean) during nodulation with the bacterial species *Bradyrhizobium japonicum* at 10 and 20 days after inoculation (50); (**G**) *Arabidopsis thaliana* and *Phaseolus vulgaris* (common bean) during infection with *Sclerotinia sclerotiorum* (93); (**H**) *A. thaliana* after infection with *Verticillium dahliae* and the *V. dahliae* mutant *aly1 aly2* (94); (**I**) *A. thaliana* infected with *Hyaloperonospora arabidopsidis* at 3, 4, and 7 dpi (95); (**J**) *Botrytis cinerea* cultivated *in vitro* (11); (**K**) *A. thaliana* infected with *B. cinerea* at 24, 48, and 72 hpi (11).

**Supplementary Figure 9. Clustering of sRNA-seq samples.** We used NMDS, PCC (Figure 3), MDS (**A** and **B**), and PCA (**C** and **D**) to estimate sample differences and similarities. Each data point represents the collapsed milRNA expression data from one sample. Blue, epiphytic fungal mycelium (MYC); green, infected epidermis without mycelium (EPI); light blue, fungal haustoria (HAU); purple, microsomes of the epidermis without haustoria (P40); orange, apoplastic extracellular vesicles (EV+); grey, apoplastic extracellular vesicles of non-infected control plants (EV-).

**Supplementary Figure 10. EV-replicate three is more similar to EV samples from infected *H. vulgare* leaves.** (**A**) Read length profiles for the three apoplastic extracellular vesicles of non-infected control plants (EV-) generated in this study. The histograms show the read counts (y-axis) for the respective read size (x-axis). (**B**) We aligned sRNA-seq reads of 31-32 bases in length to the RFAM database using MMSeqs2 (88). The stacked bar graph shows the percentage of reads identified as 5S, 5.8S, 18S, 28S, or tRNA, as indicated in the color-coded legend. Green, reads identified as derived from *H. vulgare;* blue, reads identified as derived from *B. hordei* DH14; grey, reads originating from neither *H. vulgare* nor *B. hordei;* purple, reads identified as *B. hordei* tRNA-derived. Apoplastic extracellular vesicles (EV+), apoplastic extracellular vesicles of non-infected control plants (EV-), and the three EV-replicates are shown. Total reads assigned to each sample are provided below the graph; circles visually indicate the total number of reads for comparison. (**C**) We aligned sRNA-seq reads of 31-32 bases to the *H. vulgare* 28S rDNA (3,853 bases). The graphs display the number of reads identified (y-axis) mapping to each position of the *H. vulgare* 28S rDNA (x-axis). Apoplastic extracellular vesicles (EV+), apoplastic extracellular vesicles of non-infected control plants (EV-), and the three replicates for EV-. *Hvu*-rRF0003 fragments were detected in EV+ and EV-samples, in addition to *Hvu*-rRF0004 (EV+) and *Hvu*-rRF0005 (EV-). *Hvu*-rRF0003 and *Hvu*-rRF0005 were otherwise detected only in replicate 3.

**Supplementary Figure 11. GO enrichment of putative endogenous milRNA targets.** We determined all putative targets of the sets of *B. hordei* and *H. vulgare* milRNAs via psRNAtarget (31). We used ShinyGO v0.75 (85) to calculate enriched gene ontology (GO) terms in all milRNA target sets and summarized redundant GO terms with EMBL-EBI QuickGO (https://www.ebi.ac.uk/QuickGO/) on GO version 2022-04-26 and REVIGO (86). (**A**) GO enrichment terms found in putative endogenous targets of *B. hordei* milRNAs. (**B**) GO enrichment terms found in putative endogenous targets of *H. vulgare* milRNAs. The GO terms and identifiers are indicated next to the bubble plots. Bubble size indicates fold enrichment of the term in the respective subset, fill color indicates -Log10 of the FDR-adjusted enrichment *P* value. The milRNA subsets are indicated below the bubble plots (see Figure 4 for all subsets). The icons on top of the plot were created with bioRender.com; the blue mycelium indicates *B. hordei* and the green plant barley.

**Supplementary Figure 12. PhasiRNAs are abundant in the *B. hordei* locus *BLGH_03674* encoding a pseudogenized Sgk2 kinase.**PhasiRNA-rich loci indicative of phasing in the genome of *B. hordei* were detected using ShortStack pipeline v3.8.5-1 (29), the PHASIS pipeline (33), and unitas v1.7.0 (34). The figure shows an example of a phased locus in *B. hordei*, containing the gene *BLGH_03674*, which is the partial open reading frame of a gene encoding a Sgk2 kinase. From top to bottom, the position on the scaffold, scaffold name and window, gene and transposable element annotations, and sample names are indicated. Read coverage is indicated left of each mapping profile. Red lines indicate reads mapping in sense orientation of the displayed sequence window; blue lines show reads mapping in antisense orientation. The putative trans-activating RNA (tasiRNA) is indicated with an asterisk. The figure was generated after manual inspection with Integrative Genome Viewer (IGV; (87)).

## Supplementary Data

**Supplementary File 1.** FASTA file containing all tRNA and rRNA fragments identified in this study.

**Supplementary File 2.** GFF3 file with all *H. vulgare* milRNA loci identified in this study.

**Supplementary File 3.** FASTA file with all *H. vulgare* milRNAs identified in this study.

**Supplementary File 4.** GFF3 file with all *B. hordei* milRNA loci identified in this study.

**Supplementary File 5.** FASTA file with all *B. hordei* milRNAs identified in this study.

**Supplementary Table 1.** General summary of sRNA sequencing samples, and sample description.

**Supplementary Table 2.** Read length distribution counts of the trimmed reads across all samples.

**Supplementary Table 3.** Distribution of read lengths mapped to the reference genome of *H. vulgare* (IBSCv2; (27)).

**Supplementary Table 4.** Distribution of read lengths mapped to the reference genome of *B. hordei* (28).

**Supplementary Table 5.** Reads from the samples were aligned by BLAST to the *H. vulgare* 28S rDNA; mappings were counted in 10-base windows.

**Supplementary Table 6.** Reads from the samples were aligned by BLAST to the *B. hordei* 28S rDNA; mappings were counted in 10-base windows.

**Supplementary Table 7.** Secondary structure and minimum free energy (MFE) predictions of *H. vulgare* and *B. hordei* tRNA and RNA fragments

**Supplementary Table 8.** Accession numbers of publicly available small RNA-seq datasets from barley, wheat, and Arabidopsis under biotic or abiotic stress.

**Supplementary Table 9.** Summary of unique milRNAs detected in *B. hordei* and *H. vulgare* by ShortStack, across all samples.

**Supplementary Table 10.** Counts of sRNA-seq reads mapping to the milRNA loci in *H. vulgare* identified by ShortStack.

**Supplementary Table 11.** Counts of sRNA-seq reads mapping to the milRNA loci in *B. hordei* identified by ShortStack.

**Supplementary Table 12.** GO enrichment of gene sets predicted to be targeted by *H. vulgare* milRNAs.

**Supplementary Table 13.** GO enrichment of gene sets predicted to be targeted by *B. hordei* milRNAs.

**Supplementary Table 14.** Loci and their annotation in the genome of *B. hordei* in which phasiRNAs were detected.

## References

1. Ma X, Wiedmer J, Palma-Guerrero J. 2019. Small RNA bidirectional crosstalk during the interaction between wheat and *Zymoseptoria tritici*. Front Plant Sci 10:1669. doi:10.3389/fpls.2019.01669.

2. Wong-Bajracharya J, Singan VR, Monti R, Plett KL, Ng V, Grigoriev IV, Martin FM, Anderson IC, Plett JM. 2022. The ectomycorrhizal fungus *Pisolithus microcarpus* encodes a microRNA involved in cross-kingdom gene silencing during symbiosis. Proc Natl Acad Sci USA 119. doi:10.1073/pnas.2103527119.

3. Klesen S, Hill K, Timmermans MCP. 2020. Small RNAs as plant morphogens. Curr Top Dev Biol 137:455–480. doi:10.1016/bs.ctdb.2019.11.001.

4. Chen X, Rechavi O. 2022. Plant and animal small RNA communications between cells and organisms. Nat. Rev. Mol. Cell Biol. 23:185–203. doi:10.1038/s41580-021-00425-y.

5. Bologna NG, Voinnet O. 2014. The diversity, biogenesis, and activities of endogenous silencing small RNAs in *Arabidopsis*. Annu Rev Plant Biol 65:473–503. doi:10.1146/annurev-arplant-050213-035728.

6. Lee H-C, Li L, Gu W, Xue Z, Crosthwaite SK, Pertsemlidis A, Lewis ZA, Freitag M, Selker EU, Mello CC, Liu Y. 2010. Diverse pathways generate microRNA-like RNAs and Dicer-independent small interfering RNAs in fungi. Mol. Cell 38:803–814. doi:10.1016/j.molcel.2010.04.005.

7. Komiya R. 2017. Biogenesis of diverse plant phasiRNAs involves an miRNA-trigger and Dicer-processing. J Plant Res 130:17–23. doi:10.1007/s10265-016-0878-0.

8. Lee Marzano S-Y, Neupane A, Domier L. 2018. Transcriptional and small RNA responses of the white mold fungus *Sclerotinia sclerotiorum* to infection by a virulence-attenuating hypovirus. Viruses 10. doi:10.3390/v10120713.

9. Fire A, Xu S, Montgomery MK, Kostas SA, Driver SE, Mello CC. 1998. Potent and specific genetic interference by double-stranded RNA in *Caenorhabditis elegans*. Nature 391:806–811. doi:10.1038/35888.

10. Fukudome A, Fukuhara T. 2017. Plant dicer-like proteins: double-stranded RNA-cleaving enzymes for small RNA biogenesis. J Plant Res 130:33–44. doi:10.1007/s10265-016-0877-1.

11. Weiberg A, Wang M, Lin F-M, Zhao H, Zhang Z, Kaloshian I, Huang H-D, Jin H. 2013. Fungal small RNAs suppress plant immunity by hijacking host RNA interference pathways. Science 342:118–123. doi:10.1126/science.1239705.

12. Wang M, Weiberg A, Lin F-M, Thomma BPHJ, Huang H-D, Jin H. 2016. Bidirectional cross-kingdom RNAi and fungal uptake of external RNAs confer plant protection. Nat Plants 2:16151. doi:10.1038/nplants.2016.151.

13. Cai Q, Qiao L, Wang M, He B, Lin F-M, Palmquist J, Huang H-D, Jin H. 2018. Plants send small RNAs in extracellular vesicles to fungal pathogen to silence virulence genes. Science 360:1126–1129. doi:10.1126/science.aar4142.

14. He B, Hamby R, Jin H. 2021. Plant extracellular vesicles: Trojan horses of cross-kingdom warfare. FASEB Bioadv 3:657–664. doi:10.1096/fba.2021-00040.

15. Manocha MS, Shaw M. 1964. Occurrence of lomasomes in mesophyll cells of ‘Khapli’ wheat. Nature 203:1402–1403. doi:10.1038/2031402b0.

16. Skokos D, Le Panse S, Villa I, Rousselle JC, Peronet R, David B, Namane A, Mécheri S. 2001. Mast cell-dependent B and T lymphocyte activation is mediated by the secretion of immunologically active exosomes. J. Immunol. 166:868–876. doi:10.4049/jimmunol.166.2.868.

17. Segura E, Guérin C, Hogg N, Amigorena S, Théry C. 2007. CD8^+^ dendritic cells use LFA-1 to capture MHC-peptide complexes from exosomes in vivo. J. Immunol. 179:1489–1496. doi:10.4049/jimmunol.179.3.1489.

18. Morelli AE, Larregina AT, Shufesky WJ, Sullivan MLG, Stolz DB, Papworth GD, Zahorchak AF, Logar AJ, Wang Z, Watkins SC, Falo LD, Thomson AW. 2004. Endocytosis, intracellular sorting, and processing of exosomes by dendritic cells. Blood 104:3257–3266. doi:10.1182/blood-2004-03-0824.

19. Parolini I, Federici C, Raggi C, Lugini L, Palleschi S, Milito A de, Coscia C, Iessi E, Logozzi M, Molinari A, Colone M, Tatti M, Sargiacomo M, Fais S. 2009. Microenvironmental pH is a key factor for exosome traffic in tumor cells. J. Biol. Chem. 284:34211–34222. doi:10.1074/jbc.M109.041152.

20. Woith E, Fuhrmann G, Melzig MF. 2019. Extracellular vesicles-connecting kingdoms. Int J Mol Sci 20. doi:10.3390/ijms20225695.

21. Zand Karimi H, Baldrich P, Rutter BD, Borniego L, Zajt KK, Meyers BC, Innes RW. 2022. Arabidopsis apoplastic fluid contains sRNA- and circular RNA-protein complexes that are located outside extracellular vesicles. Plant Cell 34:1863–1881. doi:10.1093/plcell/koac043.

22. Liu M, Braun U, Takamatsu S, Hambleton S, Shoukouhi P, Bisson KR, Hubbard K. 2021. Taxonomic revision of *Blumeria* based on multi-gene DNA sequences, host preferences and morphology. Mycoscience 62:143–165. doi:10.47371/mycosci.2020.12.003.

23. Kusch S, Frantzeskakis L, Thieron H, Panstruga R. 2018. Small RNAs from cereal powdery mildew pathogens may target host plant genes. Fungal Biol. 122:1050–1063. doi:10.1016/j.funbio.2018.08.008.

24. Hunt M, Banerjee S, Surana P, Liu M, Fuerst G, Mathioni S, Meyers BC, Nettleton D, Wise RP. 2019. Small RNA discovery in the interaction between barley and the powdery mildew pathogen. BMC Genomics 20:345. doi:10.1186/s12864-019-5947-z.

25. Hippe S. 1985. Ultrastructure of haustoria of *Erysiphe graminis* f. sp. hordei preserved by freeze-substitution. Protoplasma 129:52–61. doi:10.1007/BF01282305.

26. An QL, Ehlers K, Kogel KH, van Bel AJE, Huckelhoven R. 2006. Multivesicular compartments proliferate in susceptible and resistant MLA12-barley leaves in response to infection by the biotrophic powdery mildew fungus. New Phytol. 172:563–576.

27. Mascher M, Gundlach H, Himmelbach A, Beier S, Twardziok SO, Wicker T, Radchuk V, Dockter C, Hedley PE, Russell J, Bayer M, Ramsay L, Liu H, Haberer G, Zhang X-Q, Zhang Q, Barrero RA, Li L, Taudien S, Groth M, Felder M, Hastie A, Simkova H, Stankova H, Vrana J, Chan S, Munoz-Amatriain M, Ounit R, Wanamaker S, Bolser D, Colmsee C, Schmutzer T, Aliyeva-Schnorr L, Grasso S, Tanskanen J, Chailyan A, Sampath D, Heavens D, Clissold L, Cao S, Chapman B, Dai F, Han Y, Li H, Li X, Lin C, McCooke JK, Tan C, Wang P, Wang S, Yin S, Zhou G, Poland JA, Bellgard MI, Borisjuk L, Houben A, Dolezel J, Ayling S, Lonardi S, Kersey P, Langridge P, Muehlbauer GJ, Clark MD, Caccamo M, Schulman AH, Mayer KFX, Platzer M, Close TJ, Scholz U, Hansson M, Zhang G, Braumann I, Spannagl M, Li C, Waugh R, Stein N. 2017. A chromosome conformation capture ordered sequence of the barley genome. Nature 544:427–433. doi:10.1038/nature22043.

28. Frantzeskakis L, Kracher B, Kusch S, Yoshikawa-Maekawa M, Bauer S, Pedersen C, Spanu PD, Maekawa T, Schulze-Lefert P, Panstruga R. 2018. Signatures of host specialization and a recent transposable element burst in the dynamic one-speed genome of the fungal barley powdery mildew pathogen. BMC Genomics 19:27. doi:10.1186/s12864-018-4750-6.

29. Johnson NR, Yeoh JM, Coruh C, Axtell MJ. 2016. Improved placement of multi-mapping small RNAs. G3 6:2103–2111. doi:10.1534/g3.116.030452.

30. Gruber AR, Lorenz R, Bernhart SH, Neuböck R, Hofacker IL. 2008. The Vienna RNA websuite. Nucleic Acids Res 36:W70–4. doi:10.1093/nar/gkn188.

31. Dai X, Zhao PX. 2011. psRNATarget: A plant small RNA target analysis server. Nucleic Acids Res 39:W155–9. doi:10.1093/nar/gkr319.

32. Kusch S, Ahmadinejad N, Panstruga R, Kuhn H. 2014. *In silico* analysis of the core signaling proteome from the barley powdery mildew pathogen *(Blumeria graminis* f.sp. *hordei)*. BMC Genomics 15:843. doi:10.1186/1471-2164-15-843.

33. Kakrana A, Li P, Patel P, Hammond R, Anand D, Mathioni SM, Meyers BC. 2017. PHASIS : A computational suite for de novo discovery and characterization of phased, siRNA-generating loci and their miRNA triggers. bioRxiv. doi:10.1101/158832.

34. Gebert D, Hewel C, Rosenkranz D. 2017. unitas: The universal tool for annotation of small RNAs. BMC Genomics 18:644. doi:10.1186/s12864-017-4031-9.

35. Guo Q, Qu X, Jin W. 2015. PhaseTank: Genome-wide computational identification of phasiRNAs and their regulatory cascades. Bioinformatics 31:284–286. doi:10.1093/bioinformatics/btu628.

36. Li Z, Ender C, Meister G, Moore PS, Chang Y, John B. 2012. Extensive terminal and asymmetric processing of small RNAs from rRNAs, snoRNAs, snRNAs, and tRNAs. Nucleic Acids Res 40:6787–6799. doi:10.1093/nar/gks307.

37. Chen Z, Sun Y, Yang X, Wu Z, Guo K, Niu X, Wang Q, Ruan J, Bu W, Gao S. 2017. Two featured series of rRNA-derived RNA fragments (rRFs) constitute a novel class of small RNAs. PLoS One 12:e0176458. doi:10.1371/journal.pone.0176458.

38. Lee H-C, Chang S-S, Choudhary S, Aalto AP, Maiti M, Bamford DH, Liu Y. 2009. qiRNA is a new type of small interfering RNA induced by DNA damage. Nature 459:274–277. doi:10.1038/nature08041.

39. Asha S, Soniya EV. 2017. The sRNAome mining revealed existence of unique signature small RNAs derived from 5.8SrRNA from *Piper nigrum* and other plant lineages. Sci Rep 7:41052. doi:10.1038/srep41052.

40. Guan L, Grigoriev A. 2020. Age-related argonaute loading of ribosomal RNA fragments. MicroRNA 9:142–152. doi:10.2174/2211536608666190920165705.

41. Zhou X, Feng X, Mao H, Li M, Xu F, Hu K, Guang S. 2017. RdRP-synthesized antisense ribosomal siRNAs silence pre-rRNA via the nuclear RNAi pathway. Nat Struct Mol Biol 24:258–269. doi:10.1038/nsmb.3376.

42. Zhu C, Yan Q, Weng C, Hou X, Mao H, Liu D, Feng X, Guang S. 2018. Erroneous ribosomal RNAs promote the generation of antisense ribosomal siRNA. Proc Natl Acad Sci USA 115:10082–10087. doi:10.1073/pnas.1800974115.

43. Alves CS, Nogueira FTS. 2021. Plant small RNA world growing bigger: tRNA-derived fragments, longstanding players in regulatory processes. Front Mol Biosci 8:638911. doi:10.3389/fmolb.2021.638911.

44. Lalande S, Merret R, Salinas-Giegé T, Drouard L. 2020. Arabidopsis tRNA-derived fragments as potential modulators of translation. RNA Biol 17:1137–1148. doi:10.1080/15476286.2020.1722514.

45. Hackenberg M, Huang P-J, Huang C-Y, Shi B-J, Gustafson P, Langridge P. 2013. A comprehensive expression profile of microRNAs and other classes of non-coding small RNAs in barley under phosphorous-deficient and -sufficient conditions. DNA Res 20:109–125. doi:10.1093/dnares/dss037.

46. Hsieh L-C, Lin S-I, Shih AC-C, Chen J-W, Lin W-Y, Tseng C-Y, Li W-H, Chiou T-J. 2009. Uncovering small RNA-mediated responses to phosphate deficiency in Arabidopsis by deep sequencing. Plant Physiol 151:2120–2132. doi:10.1104/pp.109.147280.

47. Sun Z, Hu Y, Zhou Y, Jiang N, Hu S, Li L, Li T. 2022. tRNA-derived fragments from wheat are potentially involved in susceptibility to Fusarium head blight. BMC Plant Biol 22:3. doi:10.1186/s12870-021-03393-9.

48. Ma X, Liu C, Kong X, Liu J, Zhang S, Liang S, Luan W, Cao X. 2021. Extensive profiling of the expressions of tRNAs and tRNA-derived fragments (tRFs) reveals the complexities of tRNA and tRF populations in plants. Sci China Life Sci 64:495–511. doi:10.1007/s11427-020-1891-8.

49. Nunes CC, Gowda M, Sailsbery J, Xue M, Chen F, Brown DE, Oh Y, Mitchell TK, Dean RA. 2011. Diverse and tissue-enriched small RNAs in the plant pathogenic fungus, *Magnaporthe oryzae*. BMC Genomics 12:288. doi:10.1186/1471-2164-12-288.

50. Ren B, Wang X, Duan J, Ma J. 2019. Rhizobial tRNA-derived small RNAs are signal molecules regulating plant nodulation. Science 365:919–922. doi:10.1126/science.aav8907.

51. Martinez G, Choudury SG, Slotkin RK. 2017. tRNA-derived small RNAs target transposable element transcripts. Nucleic Acids Res 45:5142–5152. doi:10.1093/nar/gkx103.

52. Yamasaki S, Ivanov P, Hu G-F, Anderson P. 2009. Angiogenin cleaves tRNA and promotes stress-induced translational repression. J. Cell Biol. 185:35–42. doi:10.1083/jcb.200811106.

53. Thompson DM, Parker R. 2009. The RNase Rny1p cleaves tRNAs and promotes cell death during oxidative stress in *Saccharomyces cerevisiae*. J. Cell Biol. 185:43–50. doi:10.1083/jcb.200811119.

54. Singh NK, Paz E, Kutsher Y, Reuveni M, Lers A. 2020. Tomato T2 ribonuclease LE is involved in the response to pathogens. Mol Plant Pathol 21:895–906. doi:10.1111/mpp.12928.

55. Megel C, Hummel G, Lalande S, Ubrig E, Cognat V, Morelle G, Salinas-Giegé T, Duchêne A-M, Maréchal-Drouard L. 2019. Plant RNases T2, but not Dicer-like proteins, are major players of tRNA-derived fragments biogenesis. Nucleic Acids Res 47:941–952. doi:10.1093/nar/gky1156.

56. Alves CS, Vicentini R, Duarte GT, Pinoti VF, Vincentz M, Nogueira FTS. 2017. Genome-wide identification and characterization of tRNA-derived RNA fragments in land plants. Plant Mol. Biol. 93:35–48. doi:10.1007/s11103-016-0545-9.

57. Park C-J, Caddell DF, Ronald PC. 2012. Protein phosphorylation in plant immunity: Insights into the regulation of pattern recognition receptor-mediated signaling. Front Plant Sci 3:177. doi:10.3389/fpls.2012.00177.

58. Mueth NA, Ramachandran SR, Hulbert SH. 2015. Small RNAs from the wheat stripe rust fungus *(Puccinia striiformis* f.sp. *tritici)*. BMC Genomics 16:718. doi:10.1186/s12864-015-1895-4.

59. Ji H-M, Mao H-Y, Li S-J, Feng T, Zhang Z-Y, Cheng L, Luo S-J, Borkovich KA, Ouyang S-Q. 2021. *Fol*-milR1, a pathogenicity factor of *Fusarium oxysporum*, confers tomato wilt disease resistance by impairing host immune responses. New Phytol. doi:10.1111/nph.17436.

60. Horio T. 2007. Role of microtubules in tip growth of fungi. J Plant Res 120:53–60. doi:10.1007/s10265-006-0043-2.

61. Roll-Mecak A, McNally FJ. 2010. Microtubule-severing enzymes. Curr. Opin. Cell Biol. 22:96–103. doi:10.1016/j.ceb.2009.11.001.

62. Micali CO, Neumann U, Grunewald D, Panstruga R, O’Connell R. 2011. Biogenesis of a specialized plant-fungal interface during host cell internalization of Golovinomyces orontii haustoria. Cell Microbiol. 13:210–226.

63. Movahed N, Cabanillas DG, Wan J, Vali H, Laliberté J-F, Zheng H. 2019. Turnip Mosaic Virus components are released into the extracellular space by vesicles in infected leaves. Plant Physiol 180:1375–1388. doi:10.1104/pp.19.00381.

64. Liu Y, Teng C, Xia R, Meyers BC. 2020. PhasiRNAs in plants: Their biogenesis, genic sources, and roles in stress responses, development, and reproduction. Plant Cell 32:3059–3080. doi:10.1105/tpc.20.00335.

65. Spanu PD, Abbott JC, Amselem J, Burgis TA, Soanes DM, Stüber K, van Themaat EVL, Brown JKM, Butcher SA, Gurr SJ, Lebrun MH, Ridout CJ, Schulze-Lefert P, Talbot NJ, Ahmadinejad N, Ametz C, Barton GR, Benjdia M, Bidzinski P, Bindschedler LV, Both M, Brewer MT, Cadle-Davidson L, Cadle-Davidson MM, Collemare J, Cramer R, Frenkel O, Godfrey D, Harriman J, Hoede C, King BC, Klages S, Kleemann J, Knoll D, Koti PS, Kreplak J, Lopez-Ruiz FJ, Lu XL, Maekawa T, Mahanil S, Micali C, Milgroom MG, Montana G, Noir S, O’Connell RJ, Oberhaensli S, Parlange F, Pedersen C, Quesneville H, Reinhardt R, Rott M, Sacristan S, Schmidt SM, Schön M, Skamnioti P, Sommer H, Stephens A, Takahara H, Thordal-Christensen H, Vigouroux M, Weßling R, Wicker T, Panstruga R. 2010. Genome expansion and gene loss in powdery mildew fungi reveal tradeoffs in extreme parasitism. Science 330:1543–1546.

66. Kotov AA, Adashev VE, Godneeva BK, Ninova M, Shatskikh AS, Bazylev SS, Aravin AA, Olenina LV. 2019. piRNA silencing contributes to interspecies hybrid sterility and reproductive isolation in *Drosophila melanogaster*. Nucleic Acids Res 47:4255–4271. doi:10.1093/nar/gkz130.

67. Shine MB, Zhang K, Liu H, Lim G-H, Xia F, Yu K, Hunt AG, Kachroo A, Kachroo P. 2022. Phased small RNA-mediated systemic signaling in plants. Sci. Adv. 8:eabm8791. doi:10.1126/sciadv.abm8791.

68. Li L, Collier B, Spanu P. 2019. Isolation of powdery mildew haustoria from infected barley. Bio Protoc 9. doi:10.21769/BioProtoc.3299.

69. Rutter BD, Innes RW. 2017. Extracellular vesicles isolated from the leaf apoplast carry stress-response proteins. Plant Physiol 173:728–741. doi:10.1104/pp.16.01253.

70. Barberán-Soler S, Vo JM, Hogans RE, Dallas A, Johnston BH, Kazakov SA. 2018. Decreasing miRNA sequencing bias using a single adapter and circularization approach. Genome Biol. 19:105. doi:10.1186/s13059-018-1488-z.

71. Martin M. 2011. Cutadapt removes adapter sequences from high-throughput sequencing reads. EMBnet j. 17:10. doi:10.14806/ej.17.1.200.

72. Bolger AM, Lohse M, Usadel B. 2014. Trimmomatic: A flexible trimmer for Illumina sequence data. Bioinformatics 30:2114–2120. doi:10.1093/bioinformatics/btu170.

73. Shen W, Le S, Li Y, Hu F. 2016. SeqKit: A cross-platform and ultrafast toolkit for FASTA/Q file manipulation. PLoS One 11:e0163962. doi:10.1371/journal.pone.0163962.

74. Li H, Handsaker B, Wysoker A, Fennell T, Ruan J, Homer N, Marth G, Abecasis G, Durbin R. 2009. The Sequence Alignment/Map format and SAMtools. Bioinformatics 25:2078–2079. doi:10.1093/bioinformatics/btp352.

75. Quinlan AR, Hall IM. 2010. BEDTools: A flexible suite of utilities for comparing genomic features. Bioinformatics 26:841–842. doi:10.1093/bioinformatics/btq033.

76. Liao Y, Smyth GK, Shi W. 2014. featureCounts: An efficient general purpose program for assigning sequence reads to genomic features. Bioinformatics 30:923–930. doi:10.1093/bioinformatics/btt656.

77. Wickham H. 2009. ggplot2. Elegant graphics for data analysis. Use R. Springer, New York. https://ebookcentral.proquest.com/lib/kxp/detail.action?docID=511468.

78. R Core Team. 2021. R: A language and environment for statistical computing. http://www.R-project.org/.

79. Kerpedjiev P, Hammer S, Hofacker IL. 2015. Forna (force-directed RNA): Simple and effective online RNA secondary structure diagrams. Bioinformatics 31:3377–3379. doi:10.1093/bioinformatics/btv372.

80. Langfelder P, Horvath S. 2008. WGCNA: An R package for weighted correlation network analysis. BMC Bioinformatics 9:559. doi:10.1186/1471-2105-9-559.

81. Love MI, Huber W, Anders S. 2014. Moderated estimation of fold change and dispersion for RNA-seq data with DESeq2. Genome Biol. 15:550. doi:10.1186/s13059-014-0550-8.

82. Robinson MD, McCarthy DJ, Smyth GK. 2010. edgeR: A Bioconductor package for differential expression analysis of digital gene expression data. Bioinformatics 26:139–140. doi:10.1093/bioinformatics/btp616.

83. Law CW, Chen Y, Shi W, Smyth GK. 2014. voom: Precision weights unlock linear model analysis tools for RNA-seq read counts. Genome Biol. 15:R29. doi:10.1186/gb-2014-15-2-r29.

84. Lex A, Gehlenborg N, Strobelt H, Vuillemot R, Pfister H. 2014. UpSet: Visualization of intersecting sets. IEEE Trans Vis Comput Graph 20:1983–1992. doi:10.1109/TVCG.2014.2346248.

85. Ge SX, Jung D, Yao R. 2020. ShinyGO: a graphical gene-set enrichment tool for animals and plants. Bioinformatics 36:2628–2629. doi:10.1093/bioinformatics/btz931.

86. Supek F, Bošnjak M, Škunca N, Šmuc T. 2011. REVIGO summarizes and visualizes long lists of gene ontology terms. PLoS One 6:e21800. doi:10.1371/journal.pone.0021800.

87. Robinson JT, Thorvaldsdóttir H, Wenger AM, Zehir A, Mesirov JP. 2017. Variant review with the Integrative Genomics Viewer. Cancer Res 77:e31–e34. doi:10.1158/0008-5472.CAN-17-0337.

88. Steinegger M, Söding J. 2017. MMseqs2 enables sensitive protein sequence searching for the analysis of massive data sets. Nat Biotechnol 35:1026–1028. doi:10.1038/nbt.3988.

89. Deng P, Le Wang, Cui L, Feng K, Liu F, Du X, Tong W, Nie X, Ji W, Weining S. 2015. Global identification of microRNAs and their targets in barley under salinity stress. PLoS One 10:e0137990. doi:10.1371/journal.pone.0137990.

90. Wu L, Yu J, Shen Q, Huang L, Wu D, Zhang G. 2018. Identification of microRNAs in response to aluminum stress in the roots of Tibetan wild barley and cultivated barley. BMC Genomics 19:560. doi:10.1186/s12864-018-4953-x.

91. Xin M, Wang Y, Yao Y, Song N, Hu Z, Qin D, Xie C, Peng H, Ni Z, Sun Q. 2011. Identification and characterization of wheat long non-protein coding RNAs responsive to powdery mildew infection and heat stress by using microarray analysis and SBS sequencing. BMC Plant Biol 11:61. doi:10.1186/1471-2229-11-61.

92. Ragupathy R, Ravichandran S, Mahdi MSR, Huang D, Reimer E, Domaratzki M, Cloutier S. 2016. Deep sequencing of wheat sRNA transcriptome reveals distinct temporal expression pattern of miRNAs in response to heat, light and UV. Sci Rep 6:39373. doi:10.1038/srep39373.

93. Derbyshire M, Mbengue M, Barascud M, Navaud O, Raffaele S. 2019. Small RNAs from the plant pathogenic fungus *Sclerotinia sclerotiorum* highlight host candidate genes associated with quantitative disease resistance. Mol Plant Pathol 20:1279–1297. doi:10.1111/mpp.12841.

94. Zhu C, Liu J-H, Zhao J-H, Liu T, Chen Y-Y, Wang C-H, Zhang Z-H, Guo H-S, Duan C-G. 2022. A fungal effector suppresses the nuclear export of AGO1-miRNA complex to promote infection in plants. Proc Natl Acad Sci USA 119:e2114583119. doi:10.1073/pnas.2114583119.

95. Dunker F, Trutzenberg A, Rothenpieler JS, Kuhn S, Pröls R, Schreiber T, Tissier A, Kemen A, Kemen E, Hückelhoven R, Weiberg A. 2020. Oomycete small RNAs bind to the plant RNA-induced silencing complex for virulence. eLife 9:e56096. doi:10.7554/eLife.56096.

